# Deep Learning based deconvolution methods: a systematic review

**DOI:** 10.1101/2025.02.18.638595

**Authors:** Alba Lomas Redondo, Jose M. Sánchez Velázquez, Álvaro J. García Tejedor, Víctor Javier Sánchez–Arévalo

## Abstract

Within this systematic review we examine the role of Artificial Intelligence (AI) and Deep Learning (DL) in the development of cellular deconvolution tools, with an special focus on their application to the analysis of transcriptomics data from RNA sequencing. We emphasize the critical importance of high–quality reference profiles for enhancing the accuracy of the discussed deconvolution methods, which is essential to determine cellular compositions in complex biological samples. To ensure the robustness of our work, we have applied a rigorous selection process following the Preferred Reporting Items for Systematic Reviews and Meta–Analysis (PRISMA) guidelines. Through the review process, we have identified several key research gaps, highlighting the necessity for standardized methodologies and the improvement of the interpretability of the models. Overall, we present a comprehensive, up to date overview of the different methodologies, datasets, and findings associated with DL–driven deconvolution tools, paving the way for future research and emphasizing the value of collaboration between computational and biological sciences.

## Introduction

Transcriptomics data offer a detailed view of gene expression, enabling researchers to explore cellular processes with exceptional precision. Analyzing the transcriptome is essential for tracking variations in gene expression across different conditions, identifying transcriptional isoforms and splicing events, and detecting non-coding RNA (ncRNA) species to understand their roles in various biological contexts (1). Such analyses are not only crucial for understanding fundamental molecular mechanisms but also for advancing clinical research, where transcriptomics profiling plays an increasingly important role in discovering disease biomarkers, evaluating treatment efficacy, and guiding precision medicine approaches. The regulation of gene expression is central to cellular differentiation, adaptation to environmental stimuli, and response to pathological conditions. Understanding these dynamic changes provides valuable insights into essential biological processes and disease mechanisms, including cancer progression and immune responses. Additionally, epigenetic modulation and alternative splicing greatly enhance transcript diversity, influencing cell-specific functions and phenotypic outcomes.

Given the complexity of transcriptomic data, deconvolution methods have emerged as essential tools for disentangling cellular heterogeneity, particularly in bulk RNA-seq analysis. In biomedical research and therapy development, cellular deconvolution is a key tool for characterizing the cellular composition of tissues, enabling a more detailed analysis of transcriptomic profiles in bulk RNA-seq, DNA methylation, and spatial transcriptomics studies. While specific biomarkers are primarily identified through next-generation sequencing, epigenetic profiling, and proteomic analyses, deconvolution helps contextualize their expression within different cell types, aiding in disease characterization. In oncology, for instance, biomarker identification has enabled patient stratification in breast cancer (2), melanoma (3), and lung cancer (4), facilitating personalized treatments and improving clinical outcomes. Deconvolution contributes to these advances by providing insights into immune cell infiltration, tumor heterogeneity, both of which are critical factors in disease progression and therapeutic response and and neurodegenerative conditions (5).

In the context of biology, deconvolution refers to the computational process used to estimate the proportions of different cell types within a complex tissue sample based on gene expression data. This method is crucial for characterizing the cellular composition of tissues, revealing how distinct cell populations contribute to biological processes. By disentangling these mixtures, deconvolution provides valuable insights into tissue microenvironments, supporting the identification of disease– associated cells and enabling researchers to track immune infiltration in cancer or characterize changes in inflammatory and degenerative conditions. The growing relevance of deconvolution in biomarker discovery and targeted therapy development cannot be overstated.

The high time and cost demands of detailed experimental approaches, such as single-cell RNA sequencing (scRNA-seq) or fluorescence-activated cell sorting (FACS), have led researchers to explore alternative strategies for characterizing cellular composition. Among these, computational deconvolution methods have emerged as a valuable option, enabling the estimation of cell type proportions and individual gene expression profiles from bulk RNA-seq data. These approaches help maximize the information extracted from transcriptomic datasets without the need for costly single-cell or sorting-based techniques. As transcriptomic data grow in complexity and scale, Artificial Intelligence (AI) has gained traction as a powerful tool for extracting meaningful insights. In particular, Deep Learning has driven significant innovations in computational biology and biotechnology, offering more accurate and adaptable deconvolution models.

Deep Learning models, typically represented by Deep Neural Networks, are hierarchical architectures composed of multiple layers designed to learn progressively abstract representations of the original input data (6). These models aim to improve the accuracy and efficiency of traditional deconvolution algorithms by capturing intricate, non-linear relationships within large datasets. By incorporating Deep Learning, computational deconvolution techniques can overcome the limitations of classical models, offering new insights into cellular interactions and tissue composition. These innovations have revolutionized fields such as genomic analysis, medical diagnosis, drug development, and biological network modeling, with Deep Learning’s ability to identify complex patterns and handle large volumes of data being fundamental to these advancements.

This systematic review analyzes the latest tools that have already revolutionized the field, similar to how AlphaFold (7), CellProfiler (8), and DeepVariant (9) have transformed their respective areas. The primary goal of this work is to provide a comprehensive review of state-of–the–art Deep Learning–based deconvolution methods applied to transcriptomics data. This review explores recent advances in cellular deconvolution, highlighting key methodologies, applications, and ongoing challenges in the field, while also examining the strengths and limitations of these tools. The innovation of this work lies in presenting, to the best of our knowledge, the first up–to–date and comprehensive review of these tools in the context of transcriptomics data. We offer a curated selection of relevant papers that should be considered essential reading for anyone entering this field, alongside a statistical analysis of the most relevant features of each study.

The remainder of the paper is organized as follows: In Sec.2, we revisit the mathematical foundations of convolution and the traditional methods used in transcriptomics data analysis, along with the principal Deep Learning models typically employed. In sec. 3 we describe the methodology we have followed to prepare this systematic review, making special emphasis in the selection criteria and the key research questions we consider crucial in this field. We have devoted sec. 4 to present all the results that have emerged from this review, using several tables and figures to present in a comprehensible way all the pertinent data. In sec. 5 we make a detailed discussion on the meaning of the presented results and a thoughtful reasoning of the field challenges and possible future developments. Finally, in sec. 6 we highlight the main conclusions of this review, synthesizing the main challenges we have encountered.

## Background

### Traditional deconvolution methods

In Mathematics, convolution is a well defined operator that compose any two functions into a new one, through the superposition of the former ones. Deconvolution is rigorously defined in functional analysis as the inverse operator of convolution, so given a convolved function and the filtering function used one can uniquely reconstruct the original signal.

Tissues are made up of a mixture of different cell types, and traditional deconvolution tools work under the assumption that the bulk sample is essentially described as a linear combination of gene expression from its various constituent cell types, where the coefficients of the model represent the proportions of each cell type we want to determine. This assumption can be described by the equation

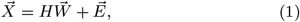

where 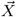 is a one–dimensional vector representing the expression of a single bulk sample that needs to be deconvolved, *H* is the gene signature matrix, 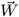 encodes the cell proportions, and 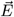 is a noise term that captures the variability and uncertainty in the observed measurements. Typically, the elements of 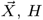 and 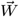 are non–negative and the cell proportions must add up to unity. When deconvolving more than one bulk sample, these variables become matrices instead of vectors.

Traditional deconvolution tools are based on probabilistic models that rely on prior knowledge about the nature of the samples. These tools utilize an input matrix or vector, *X*_*m*×*n*_, that represents bulk RNA or DNA methylation data, where *m* is the number of genes or CpG (regions where a cytosine is followed by a guanine in the DNA sequence) sites and *n* denotes the number of samples. Cellular deconvolution methods are classified into two different categories based on the level of prior knowledge they require: reference–based methods (supervised methods) and reference–free methods (unsupervised methods).

The algorithms belonging to the first category, i.e., supervised methods, require signature matrices, *H*_*m*×*k*_, corresponding to gene expression or DNA methylation of specific cell types. Here, *m* represents the number of marker genes and *k* the number of cell types. These matrices are obtained either from epigenomic or transcriptomics data of samples similar to those that wants to be deconvolved, typically using techniques like flow cytometry or FACS. The effectiveness of these methods is strongly influenced by the quality of the reference profiles. Some examples of available tools are: CIBERSORT (10), ESTIMATE (11) or EPIC (12).

On the other hand, unsupervised methods do not require predefined signature matrices, although some of them may demand prior knowledge of the specific marker genes present in the samples. In these tools, both the signature matrix and the cell proportions, *W*_*k*×*n*_, are unknown, and the goal is to estimate both matrices so they maximize the similarity between *H · W* ^*T*^ and the matrix *X*_*m*×*n*_. Some of the available tools implementing these unsupervised methods are UNDO (13), CDSeq (14) or TOAST (15).

Although these methods are widely used, they face several challenges, such as the dependence on predefined gene signatures and the assumptions on prior sample composition, depending on each model. For example, the ABSOLUTE tool (16) assumes that a cancer tissue sample is composed of a given proportion of cancer cells, *α* and a proportion of healthy cells, (1 − *α*). This assumption restricts its ability to analyse all biological variations, especially the one between stroma and tumoral cells, which is essential for predicting malignant cell proportions accurately. Furthermore, although these methods may show promising results in benchmark tests, it is still unclear how accurate they are when applied to external samples and compared to each other.

### Deep Learning Models

In recent years, the application of Deep Learning in medicine and biology has significantly increased, primarily due to its numerous advantages. This trend is evident in the growing number of scientific works published in specialized health science databases such as PubMed. As it is illustrated in Fig.1, the search results for the term *Deep Learning* in PubMed as of February 3, 2025, indicate a notable increase in the publications volume related to this topic since 2017. As a direct consequence, the use of DL methods in medicine has become widespread, with Artificial Intelligence models frequently found in both biomedical research and clinical settings (17).

**Fig. 1:**
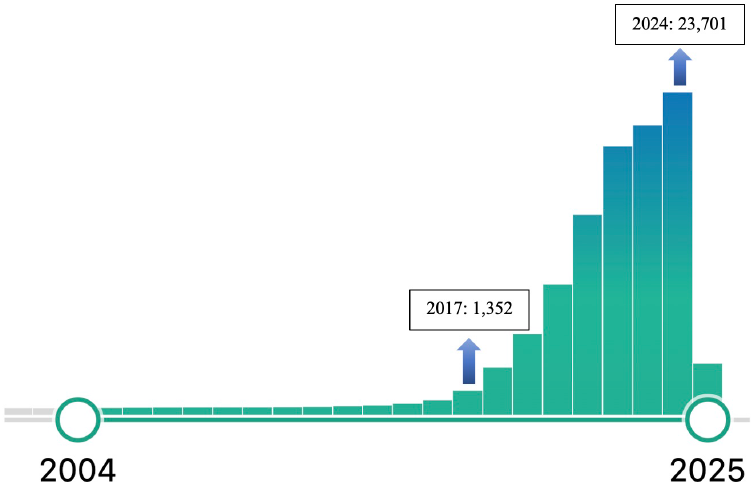
Annual publication trends of *Deep Learning* topic in PubMed.

Deep Learning currently encompasses a wide variety of Deep Neural Network models specifically tailored for different applications. Convolutional Neural Networks (CNNs) excel in image analysis, while Recurrent Neural Networks (RNN) are specially adapted for dealing with sequential data analysis. Autoencoders are used for tasks such as denoising and anomaly identification, and Generative Adversarial Networks (GANs) are employed for image generation. Additionally, Transformers have gained prominence in Natural Language Processing (NLP) for text analysis. Although each model is originally designed for specific input data types, they can be modified and combined to tackle more complex problems, leading to the creation of hybrid models (18), (19), (20). However, only a select few of these models have been applied to cellular deconvolution, as we discuss in detail in this review. Let us give a more in detail description of these models.

### Multilayer Perceptron (MLP)

This popular architecture was first introduced by Hinton (21). MLPs are composed of multiple layers of neurons –perceptrons– hierarchically arranged. The simplest form of a MLP consists of three layers: the input layer, the output layer and a single hidden layer. In contrast, deep MLPs have more than two hidden layers. Each neuron in a layer receives the same input and can apply an activation function to process the information before passing the data to the next layer. This activation function is the responsible of introducing non–linearities into the model, enabling then the MLP to capture and model complex, non–linear relationships between input and output variables. This hierarchical design allows the network to learn simple features and patterns in the initial layers while progressively identifying more abstract and complex features with each deeper layer. MLPs are particularly effective for modelling intricate relationships for a wide range of applications.

### Convolutional Neural Network (CNN)

This kind of network composes the backbone for advanced image analysis and visual recognition. Inspired by the functioning of the human visual system, CNNs process images through multiple layers, initially identifying simple features such as edges and colours, and subsequently combining these features into more complex patterns like shapes and objects. The pioneering architecture of CNNs was firstly proposed by LeCun *et al* (22), marking the beginning of its practical applications. Unlike MLPs, CNNs employ a mathematical convolution operator to analyse each pixel of the image in their corresponding neighbourhood, using a set of filters that capture specific features such as edges or textures. The selected filter slides across the image, performing a weighted sum of the surrounding pixel values using the defined filter weights. The extracted features in the early layers are typically basic or elementary; in order to capture more complex patterns, multiple sequential convolutional layers are stacked. This structure enables the network to identify increasingly abstract and sophisticated features. Additionally, CNNs reduce the data dimensionality through pooling layers while preserving relevant information, which is crucial for efficient image analysis and classification.

### Autoencoder

An autoencoder is a Deep Learning architecture specifically designed to reduce the dimensionality of data through a process of encoding and decoding. It was firstly proposed by Hinton in the 1980s (23). Although autoencoders have characteristics of both supervised and unsupervised learning, they are typically classified as unsupervised models. At their core, they consist of three key components: the encoder, which compresses the input data into a more compact, lower–dimensional representation; the bottleneck, where this compressed data is stored; and the decoder, which reconstructs the original input from the encoded form. The different layers within each of autoencoder components can vary in type, including convolutional or MLP structures, offering great flexibility in their design. Autoencoders are useful in a wide variety of applications, such as dimensionality reduction, data denoising, or anomaly detection, making them a powerful tool in data processing.

### Generative Adversarial Network

Generative adversarial networks (GANs) belong to the generative class of models. GANs consist of two neural models, the generator and the discriminator, which work together in an adversarial training process (24). The architecture of GANs aims to learn and imitate a given data distribution. The generator is responsible for producing synthetic instances of the input data, while the discriminator evaluates these instances andb decides whether they are similar enough to the input data or not. The discriminator assigns a probability of authenticity to each instance, indicating whether it is from the input distribution or synthetic. Through repeated iterations of this process, the generator learns to create synthetic data that better resembles the input distribution.

### Methodology

A systematic literature review, SLR, is a comprehensive methodological approach that identifies, evaluates and synthesizes research within a specific scientific domain or addresses defined research questions. This methodology follows clear, reproducible protocols to ensure thoroughness, reduce bias, and maintain academic rigor through transparent documentation of search strategies, inclusion criteria, and analytical methods.

Following established protocols (25), our review implements a systematic four–step process to evaluate the literature on cellular deconvolution using DL techniques. We begin by outlining the motivation behind this review, followed by a detailed examination of the selected topic to frame the research questions guiding it. These questions should be specific, designed to identify gaps in the literature, assess their relevance to the scientific community, and comprehensively address the design and evaluation of deconvolution tools.

Next, we conduct a systematic search and screening of articles based on the Preferred Reporting Items for Systematic Reviews and Meta–Analysis (PRISMA) guidelines (26; 27; 28), using their 2020 flow diagram^1^for new systematic reviews. PRISMA ensures a structured search strategy with clearly justified inclusion and exclusion criteria to focus on the most relevant studies. Each selected paper undergoes an in–depth analysis to address the initial research questions, critically evaluating the quality and methodologies employed to gain a comprehensive understanding of the field.

The final step involves synthesizing the analyzed data and presenting the findings in an organized and coherent manner. Our synthesis highlights recent advancements in cellular deconvolution using Deep Learning while identifying potential directions for future research. This structured approach ensures rigor, transparency, and replicability, enhancing the existing body of knowledge. Fig. 2 provides a flowchart summarizing these stages, serving as a roadmap for the systematic literature review process.

**Fig. 2:**
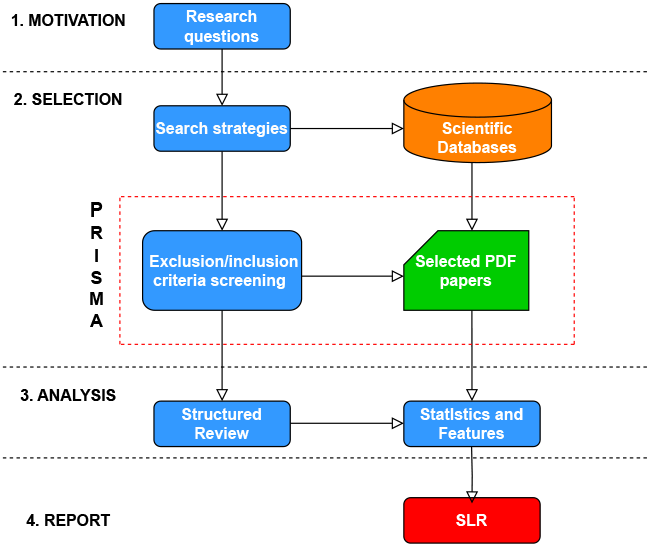
Schematic flowchart for the PRISMA approach to systematic reviewing the pertinent literature on a given topic.

### Motivation

Bulk RNA–seq data from tissue samples represents an aggregate signal derived from a heterogeneous mixture of cell types, posing significant challenges in disentangling the contributions of individual cell populations. This review examines the application of Deep Learning models for cellular deconvolution of RNA–seq data, driven by the need for more robust and reproducible techniques. Despite notable advancements in recent years, the use of Deep Learning in cellular deconvolution remains an evolving field, with several open questions and challenges that this manuscript seeks to address.

By addressing the research questions stated below in Table 1, this review offers a comprehensive overview of the current state of the art, highlighting the role of Artificial Intelligence in advancing cellular deconvolution, and identifying critical areas where further research and development are still needed. This work will not only contribute to the scientific community by summarizing existing knowledge but also guiding towards future research with potential impactful advancements in the field.

**Table 1.** Research questions guiding our systematic review.

|  |
| --- |
| **RQ 1: Datasets used by deconvolution models. What are their characteristics and availability?** |
| Finding specific transcriptomics datasets in specialized repositories can be complex. The performance of cellular deconvolution models heavily depends on their training data, requiring careful selection. Compiling and characterizing these datasets by origin, quality, and availability simplify data selection for researchers. |
| **RQ 2: Preprocessing techniques for transcriptomics data. Is there a standard procedure?** |
| scRNA-seq and RNA-seq data are often noisy and highly-dimensional, negatively impacting model performance. Preprocessing these data is crucial to ensure quality and reliability for subsequent analyses. Understanding the methods used in various studies can guide researchers in handling such data effectively. Understanding the different procedures for pseudobulk formation is also capital. |
| **RQ 3: Targeted cell types and tissues in deconvolution models. What is the purpose of each model?** |
| Deconvolution models often target specific puposes or clinical targets. Studying the final objectives of these tools, reveals the current research trends and highlights gaps where some cell types or conditions are underexplored providing guides for future research efforts addressing both well-studied and neglected areas. |
| **RQ 4: Predominant DL architectures: design, training and optimization pipelines. How are these models structured?** |
| Analyzing predominant AI-based deconvolution modelos, their design along with the full training and evaluation pipeline—including preprocessing and metrics highlights best practices that enhance reliability, reproducibility, and model comparability. |
| **RQ 5: Benchmarking. How do DL models compare to other methods?** |
| Comparing DL techniques to SOTA methods in cellular deconvolution reveals their strengths and weaknesses. This analysis helps researchers choose the most suitable techniques for their needs. It also highlights advancements and areas for future development, driving progress in the field. |
| **RQ 6: Challenges and limitations of DL in cellular deconvolution. What are the emerging trends and future directions?** |
| Identifying the main challenges and limitations of AI approaches in cellular deconvolution, such as technical variability and model interpretability, is crucial for improving these methods. Understanding emerging trends and future directions to guide research towards overcoming the challenges. |

### Search Strategy

At this stage, it is crucial to define the keywords and provide a rationale for the queries executed across multiple scientific repositories and databases. Both biological and computational disciplines must be adequately represented.

From a biological perspective, the keywords reflect both the target data and the processes analyzed. Specifically, the selected terms are: (1) cellular deconvolution, (2) deconvolution, (3) bulk RNA, and (4) RNA–seq. In contrast, the computational perspective focuses on articles exploring artificial intelligence, particularly deep neural networks (DNN), with the following keywords: (5) deep learning, (6) neural networks, and (7) artificial intelligence.

Logical operators were used to combine these terms and ensure comprehensive query coverage. To retrieve papers relevant to both fields, the biological and computational terms were linked using the AND operator. The biological keywords are divided into two groups —biological processes (1 OR 2) and target data (3 OR 4)— which are required simultaneously through the AND operator. Computational keywords were combined using the OR operator. Consequently, the final query structure was defined as: (1 OR 2) AND (3 OR 4) AND (5 OR 6 OR 7).

This search strategy was applied to six major databases: Web of Science (WOS), PubMed, ScienceDirect, Scopus, IEEE Xplore, and SpringerLink. It is worth noting that MEDLINE is indexed in PubMed, so articles stored in MEDLINE are also included in PubMed searches. The complete search process, covering all publications available up to February 3, 2025, is illustrated in Fig.3.

**Fig. 3:**
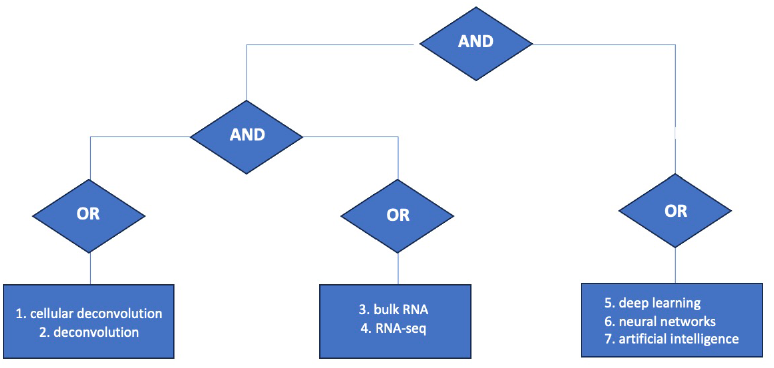
Flow diagram of the connectors employed in the PRISMA search.

### PRISMA

To ensure the selection of high–quality papers, a set of criteria was established by a multidisciplinary team comprising computer scientists, biomedical engineers, and biotechnologists. In this phase, the PRISMA guidelines were applied, as described in the following text and illustrated in Fig.4.

### Identification

A comprehensive search of all relevant literature from electronic databases was conducted according to the search strategy. There is no restriction on the publication date of the search results. Duplicate papers were discarded to ensure the uniqueness of the collected records. Also, non–journal papers (academic books, book chapters, posters, proceedings, abstracts and dissertations), reviews and non–peer reviewed papers (arXiv papers) were discarded to ensure the relevance of the records collected.

### Screening

In the initial evaluation stage, the titles and abstracts of the collected literature were reviewed to assess their relevance. This review was conducted by two researchers with contrasting backgrounds: a computer scientist and a bioscientist. This step ensures that the perspective of both fields is covered. To this end, two inclusion criteria are used:

1. Papers that explicitly mention in their title or abstract the use of artificial intelligence and deconvolution tools applied to transcriptomics data.
2. Studies that do not explicitly specify in their title or abstract the need for additional data types, other than RNA–seq or scRNA–seq, for the deconvolution tool.

Studies that do not meet the inclusion criteria are excluded, narrowing down the pool of potential works and ensuring that only high–quality papers were retained for further review. Papers identified as relevant in this step were pursued for full–text retrieval and more detailed assessment. A final selection of papers was made using stricter criteria, known as exclusion criteria. All papers fulfilling any of the criteria specified in Table 2 are excluded from the analysis.

**Table 2.** PRISMA–based exclusion criteria for a systematic literature review.

| Exclusion criterion | Reason of exclusion |
| --- | --- |
| Papers whose deconvolution tool requires other types of data. | Mandatory use of data other than RNA-seq and scRNA-seq is out of the scope of this review (bioscience criteria). |
| Papers that do not use DL techniques. | Some authors classify ML models within the DL category; however, ML models fall outside the scope of this review (computers sciences criteria). |
| Papers using DL models to process transcriptomics data but not to perform deconvolution. | Using existing deconvolution methods to subsequently design a DL-based method for other purposes is out of the scope of this review (bioscience criteria). |
| Papers that use existing DL-based deconvolution tools applied to a specific use case. | Out of scope (bioscience criteria). |

Although all studies contained the specified search terms, some only used them as references to other works or mentioned them without further development. Studies that could not be downloaded were discarded.

### Inclusion

Studies that meet all inclusion criteria and pass the exclusion filter are included in the systematic review and meta–analysis. Comprehensive data from these sources are extracted and compiled, forming a robust foundation for addressing the proposed research questions.

### Analysis

This phase offers a structured overview of the current research landscape, emphasizing key findings, identifying limitations, and proposing future directions. The main steps include:

1. **Data Extraction and Synthesis**: Collect and synthesize information from the selected studies, including objectives, methodologies, datasets, DL techniques, preprocessing steps, evaluation metrics, and outcomes.
2. **Comparative Analysis**: Evaluate the effectiveness, advantages, and limitations of different DL techniques to identify the most promising approaches and opportunities for further research.
3. **Quality Assessment**: Evaluate the scientific rigor and reliability of the included studies using predefined quality criteria.
4. **Gap Identification and Future Directions**: Highlight research gaps and propose future pathways to advance cellular deconvolution techniques using DL.

## Results

### Selection and screening process

The search was conducted on February 03, 2025, collecting articles published up to that date. This resulted in the acquisition of 171 articles, which were subsequently downloaded to the Zotero platform for a more detailed analysis. The distribution of articles across the databases was as follows: Scopus 28, PubMed 23, WOS 37, ScienceDirect 9, IEEexplore 2, SpringerLink 72. Of the initial 171 papers, 42 were duplicate items, and 23 were classified as reviews, nonjournal, or nonpeer–reviewed articles. The remaining papers (*n* = 106) were included in the screening phase. In this phase, 50 articles were eliminated because they did not meet the criterion of being a Deep Learning study developed to deconvolve transcriptomics data. Therefore, 56 studies were fully analyzed, of which 41 met the exclusion criteria for the review. Thus, a total of 13 papers formed the final selection. Fig. 4 shows these results in detail.

**Fig. 4:**
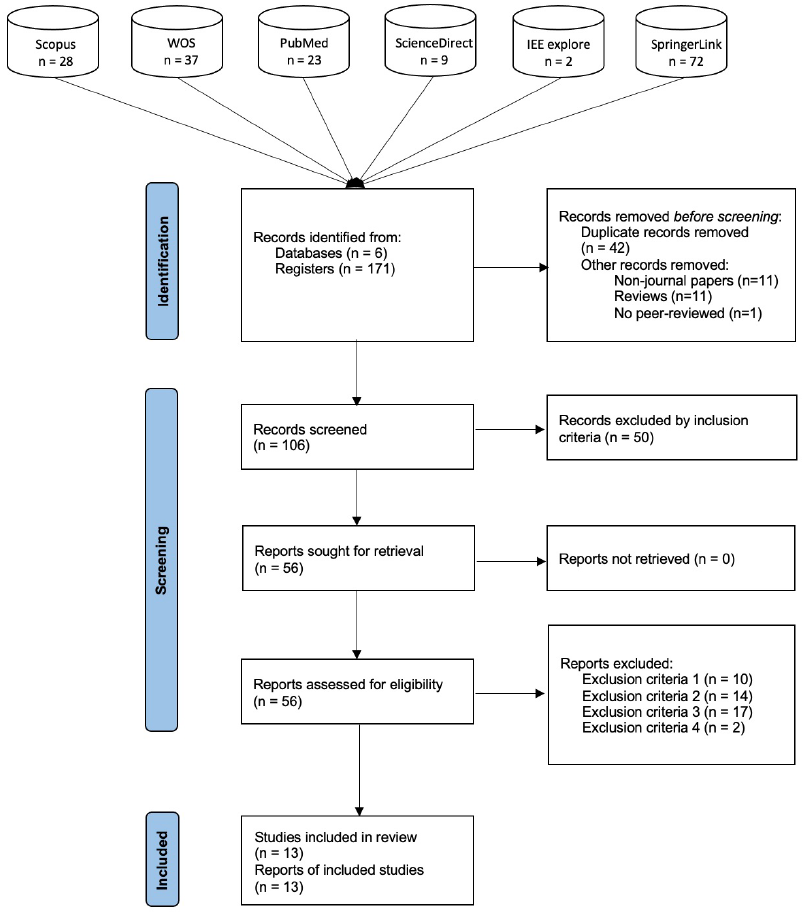
Flow diagram following the PRISMA guidelines, showing the identification, screening, eligibility assessment, and inclusion of studies in the SLR.

### Summary of papers

Fig. 5 shows the distribution of the number of DL–based deconvolution tools published by year since 2019, when the first paper in this review was published. An increasing trend in publications is evident, with a possible peak observed in 2024. This trend reflects the growing interest in the field; notably, despite 2025 having just begun, a new publication introducing such tools has already been released. Fig.5 also highlights that the development of these tools is still in its early stages, indicating a promising and emerging research niche.

**Fig. 5:**
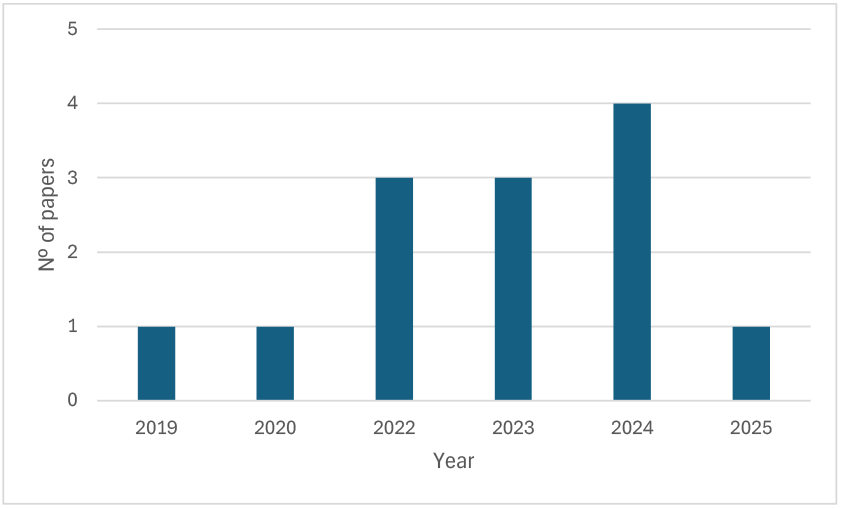
Annual number of papers on DL–based deconvolution tools per year graph.

The following paragraphs provide a general summary of the key highlights from the selected papers. The most relevant information has been compiled in Tables 3 and 4. Table 3 outlines the biological targets of each study and the approaches used for transcriptomics data processing. Meanwhile, Table 4 focuses on critical aspects of the DL models, including evaluation metrics, selected hyperparameters, and implementation frameworks.

**Table 3.**
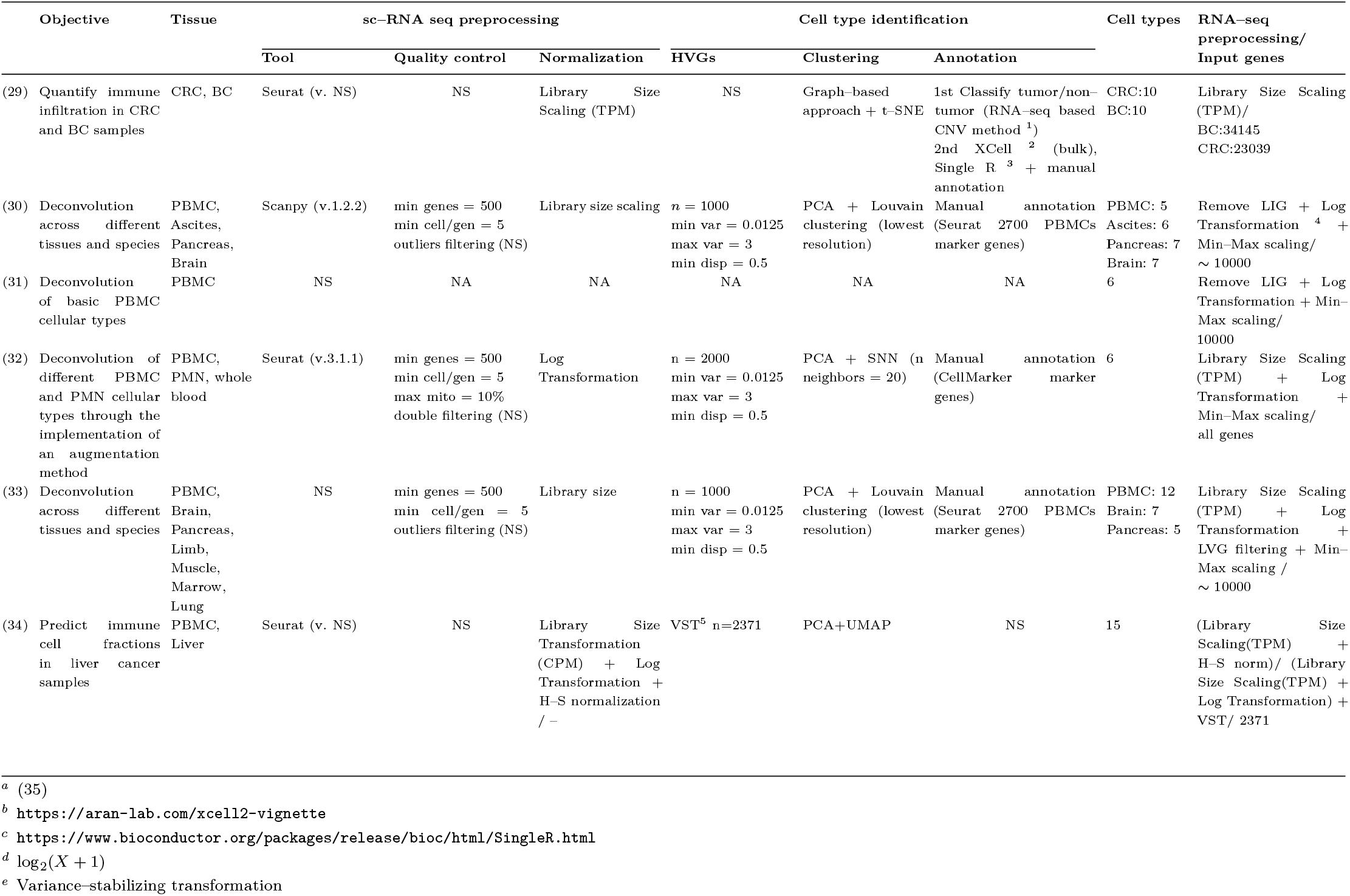
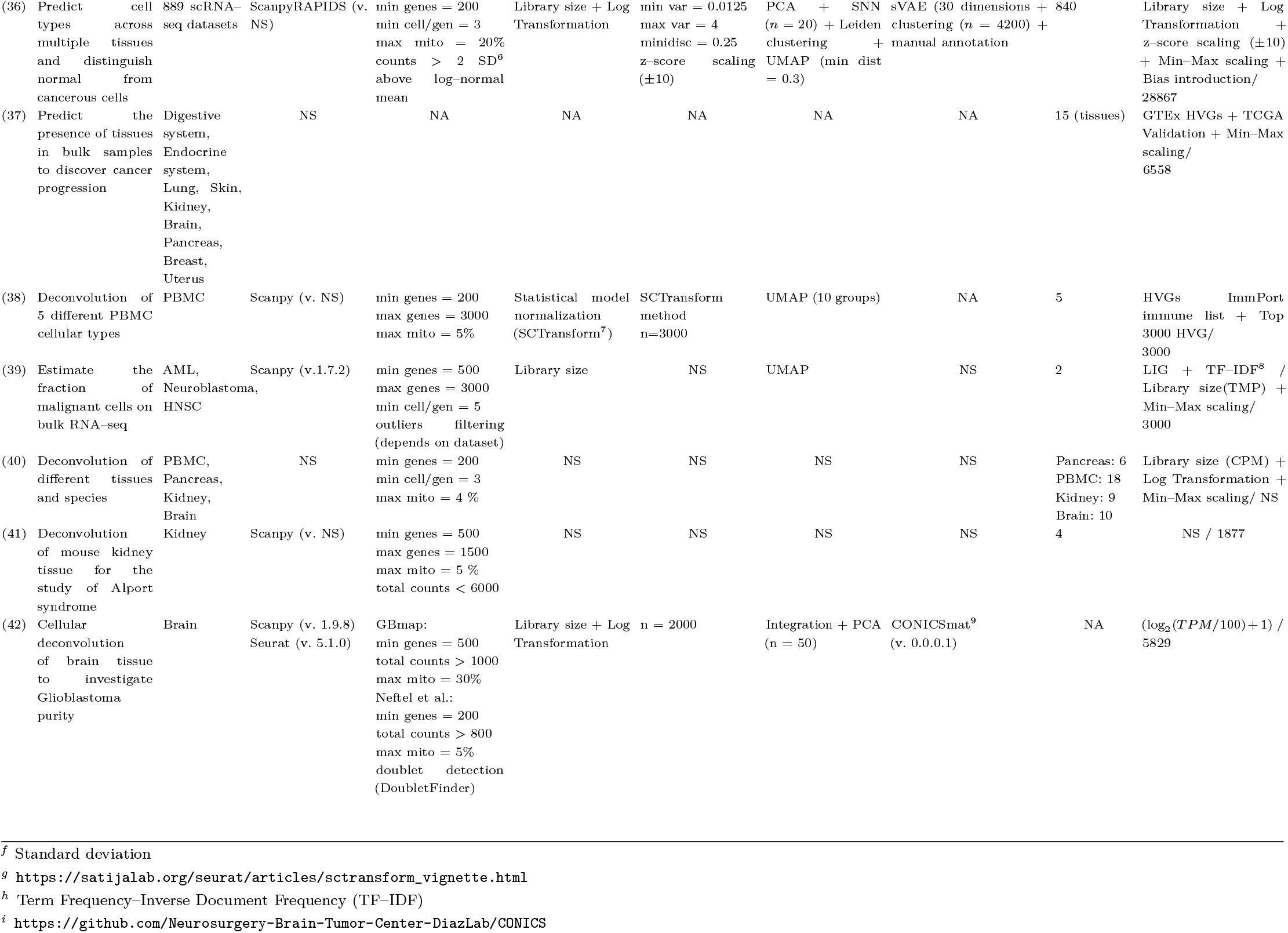
Biological Attributes for Deconvolution Techniques Summarized in This Review. The scRNA–seq preprocessing section is organized into quality control. and normalization steps. In the Quality Control section, several parameters are defined: min genes indicates the minimum number of genes required for a cell to pass filtering and avoid removal due to sequencing errors; max genes specifies the threshold above which a cell is flagged as a potential doublet; min cell/gen is the minimum number of cells that must express a gene for it to be retained as relevant; and max mito denotes the maximum allowable percentage of mitochondrial genes, beyond which a cell is excluded, indicating potential cell damage or death. The Cell Type Identification section details the identification of highly variable genes (HVGs), clustering of cells into types, and cell annotation. In identifying HVGs, *n* represents the number of selected HVGs, min var is the minimum variance required for a gene to qualify as an HVG, max var is the maximum gene variance allowed, and min disp is the minimum dispersion threshold. The Cell Types section lists the number of cell types being deconvolved, which aligns with the model output. The RNA–seq processing section describes data handling, typically involving normalization and filtering of low–information genes (LIG), which are defined as genes with zero expression or with expression variance below 0.1. Additionally, the number of genes input into the model is specified. ‘NS’ stands for ‘Not Specified’, indicating missing information in the main corpus, and ‘NA’ stands for ‘Not Applicable’, indicating data or categories that do not apply to the tool or analysis.

**Table 4.**
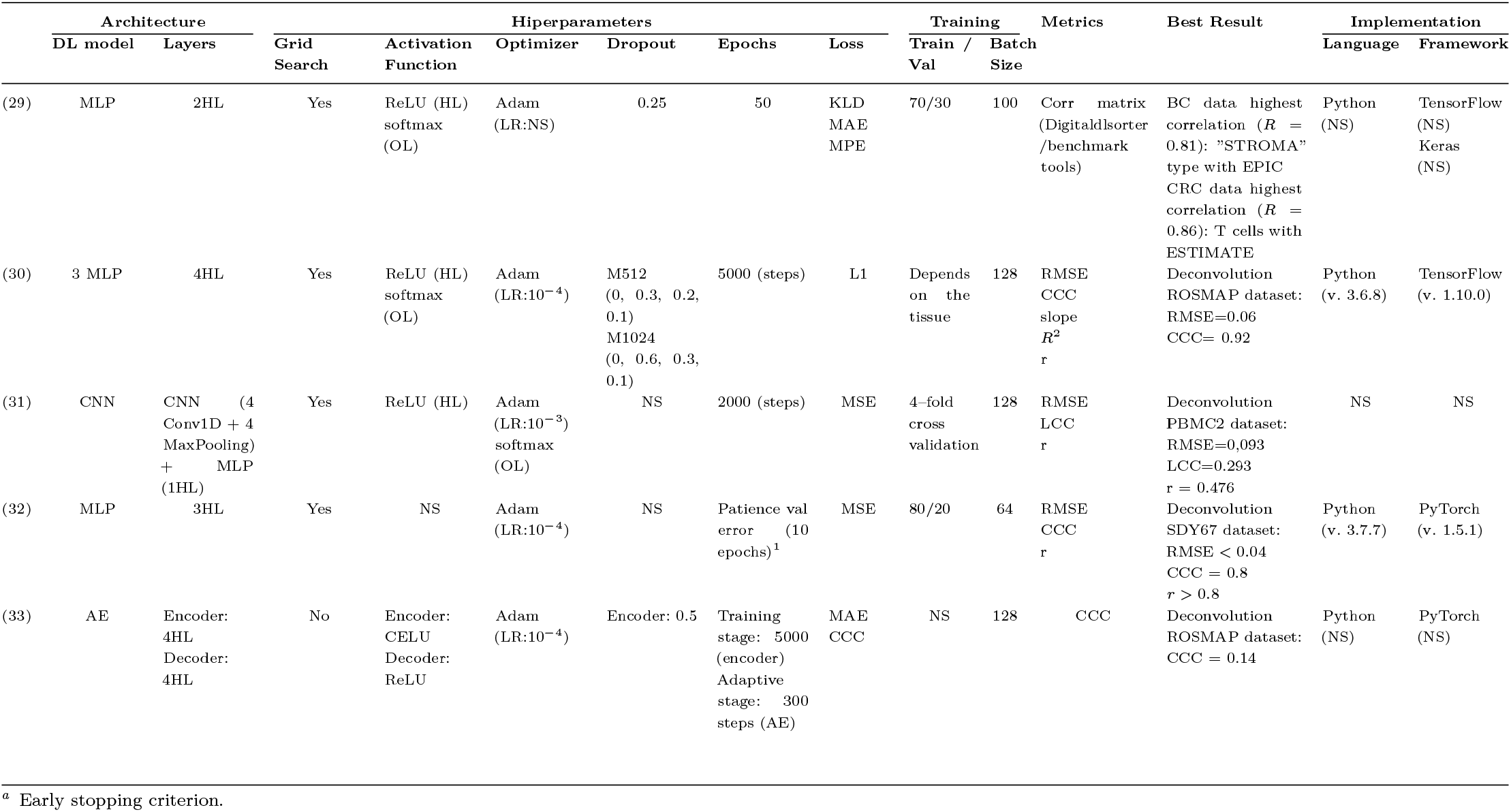
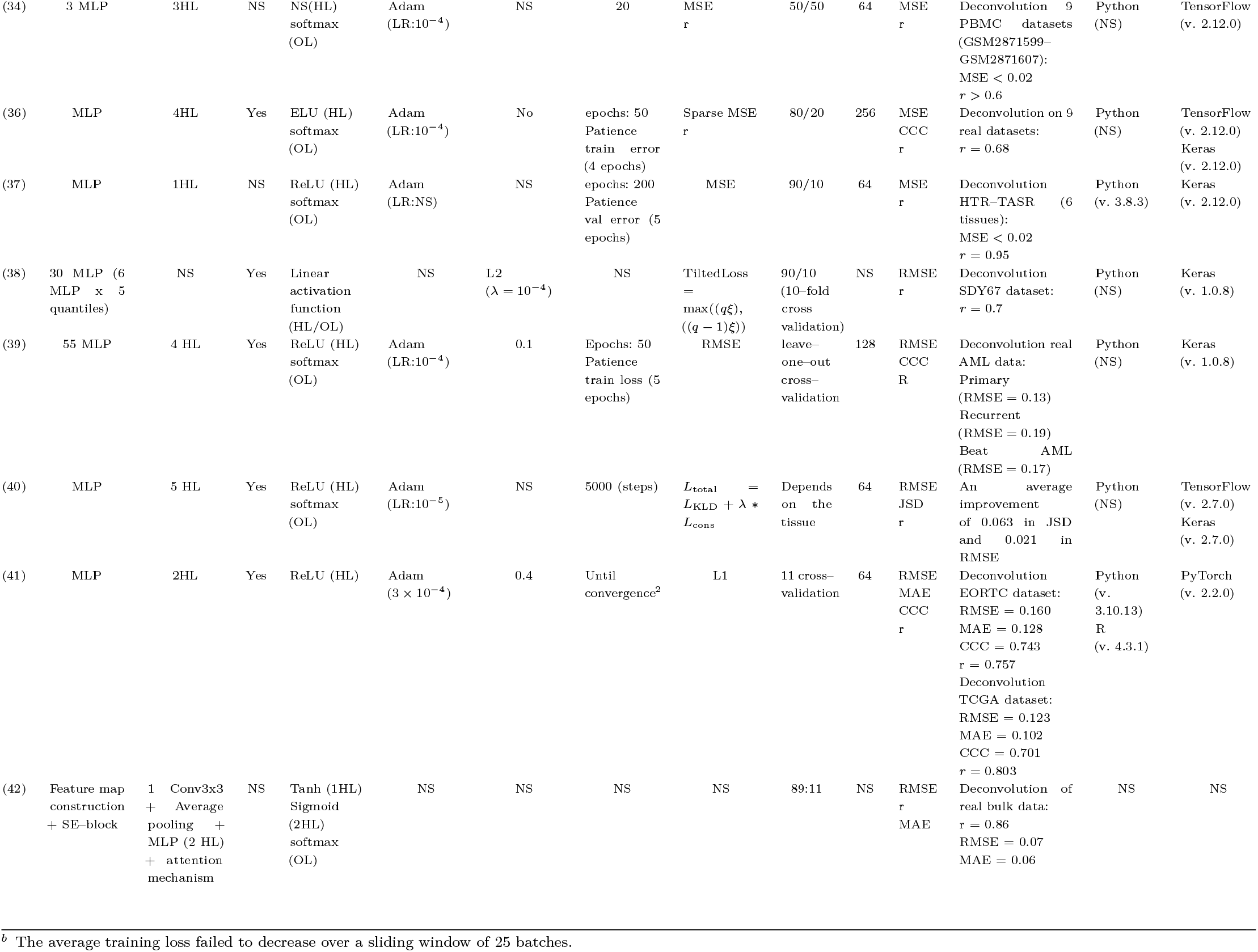
Summary of Deep Learning architectures, hyperparameters, training setups, and performance metrics used in deconvolution models. Each row presents details of a specific study, including its architecture, key hyperparameters (e.g., activation functions, optimizers, dropout rates), and training configurations (e.g., loss functions, training–validation split, batch size). Grid Search section shows whether a grid search has been performed to optimize the model architecture. Loss and Metrics section specifies the chosen functions for training and evaluating the models respectively. Best Result column contains the best achieved results of each tool in its benchmarking. The Implementation column specifies the programming language and Deep Learning framework used for each model.

1. Torroja and Sánchez–Cabo, 2019 introduced a novel method, Digitaldlsorter, for deconvoluting bulk RNA-Seq based on DL models. Digitaldlsorter is focused on the deconvolution of the immunogenic environment in breast (BC) and colorrectal (CRC) cancer samples (29).
2. Menden et al., 2020 reported Scaden, a model composed of three MLPs designed for the deconvolution of bulk transcriptomic data across various tissues, including the pancreas, brain, ovarian tissue, and PBMCs, in different species and sequencing technologies (30).
3. Zhendong et al., 2022 described a deconvolution model based on a CNN, Autoptcr, focusing on the study of the main cell types found in PBMC samples (31).
4. Lin et al., 2022 propose a pipeline called DAISM–DNN, which employs a novel in silico data augmentation strategy (DAISM) together with a MLP model to deconvolute samples from PBMC data (32).
5. Chen et al., 2022, pioneered the use of an autoencoder model for the deconvolution and generation of Gene Expression Profiles from RNA–seq samples. Their approach, TAPE, demonstrates the ability to deconvolve samples across various species and multiple organs or tissues (33).
6. Chiu et al., 2023, designed a deconvolution pipeline (HASCAD) that combines Harmony–Symphony correction with a three MLP structure of three hidden layers. This model is applied to investigate the impacts of immune cell heterogeneity on therapeutic effects, especially in cancer immunotherapy (34).
7. Charytonowicz et al., 2023, described UCDBase, an MLP–based deconvolution method with an extension called UCDSelect that uses reference data to enhance predictions. UCD study ischemic kidney injury, cancer subtypes, and tumor microenvironments by deconvolving 840 different cell types (36).
8. Yan et al., 2023, introduced an MLP–based DL model for tissue deconvolution of ctcRNA. This model is applied to study 15 different tissues and the migration of ctcRNA in a cancer patient with metastatic tumors to explore the early detection of metastasis (37).
9. Jin et al., 2024 designed NNICE, a novel method that integrates quantile regression with DL techniques to estimate cell type abundance and its variability from bulk RNA-seq data, providing an efficient and reliable solution for PBMC samples deconvolution (38).
10. Huang et al., 2024, developed DeepDecon, an iterative DL– based deconvolution tool designed to estimate the fraction of malignant cells in bulk RNA–seq samples. DeepDecon is intended for use in cancer detection, recurrence monitoring, and prognosis (39).
11. Khatri et al., 2024, reported the use of two DL networks with simultaneous consistency regularization of the target and training domains known as DISSECT. It is capable of deconvolving a variety of tissues and is easily adapted to other biomedical data types (40).
12. Sun et al. (2024) investigated changes in macrophage and T lymphocyte concentrations in kidney tissue from both healthy mice and those affected by Alport syndrome, employing a hybrid model (CONVdecov) incorporating convolutional layers and attention mechanisms. (41).
13. Morgan et al. (2025) designed a tool, GBMPurity, to estimate tumor purity in glioblastomas from human tissue, using an interpretable DL model based on a simple MLP (42).

### Datasets used by deconvolution models

This section examines the datasets utilized in the reviewed studies, focusing on their specific characteristics. The goal is to clarify the types of data employed in developing deconvolution tools with DL, their biological origins, and the sources from which they were acquired. The first part of the analysis identifies the various data types used in the development of the deconvolution tools covered in this review. Fig. 6 illustrates the diversity of genomic data and their distribution, highlighting the number of studies in which each type is featured. Each paper, regardless of whether it employs multiple datasets of the same type or just one, is counted equally. For this section, UCDBase tool has been excluded due to its extensive dataset collection (over 1000 datasets), which makes detailed analysis unfeasible.

**Fig. 6:**
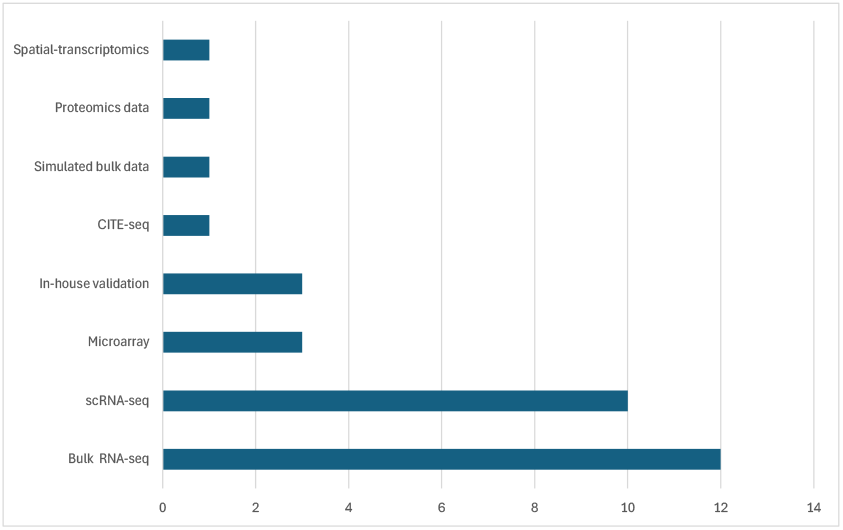
Distribution of Genomic Data Types Used in Deconvolution Studies. Simulated bulk data category refers to studies where the data are not simulated, but instead, pre–existing simulated data are used, and In–house validation category refers to studies where the data are generated or collected internally by a team or laboratory to validate and assess the model performance.

As we show, all studies (*n* = 12) use bulk RNA–seq data, in consonance with the search criteria described in the methodology section. In most of the cases (*n* = 10), scRNA–seq data is also required to generate pseudobulk data. Data from microarray technologies appears in 23% of the studies, while other types of data (spatial transcriptomics and proteomics) are used less frequently. It is also notable that three of the studies use data generated by the authors themselves.

Focusing on the predominant data types (bulk RNA–seq and scRNA–seq), the next questions concern the origin of these datasets, specifically the species and tissue types targeted for deconvolution. In Fig. 7 we show the predominance of *Homo sapiens* data compared to *Mus musculus* data, with the former one used in all studies and the latter appearing in only four out of twelve. Meanwhile, in Fig. 8 we illustrate the distribution of tissue origins for the datasets, emphasizing the predominance of data from PBMC samples, followed by cancer tissue studies, as well as pancreatic and brain tissue.

**Fig. 7:**
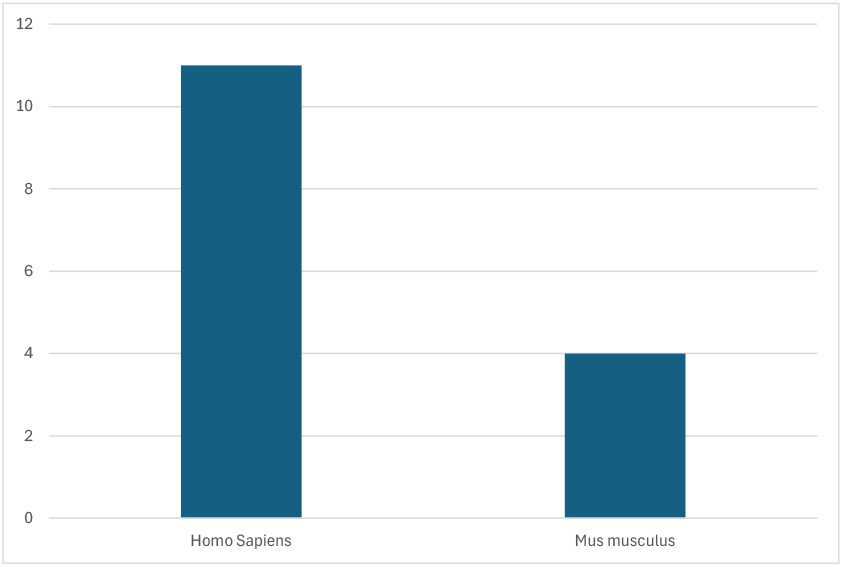
Species Distribution in Datasets Used for Deconvolution.

**Fig. 8:**
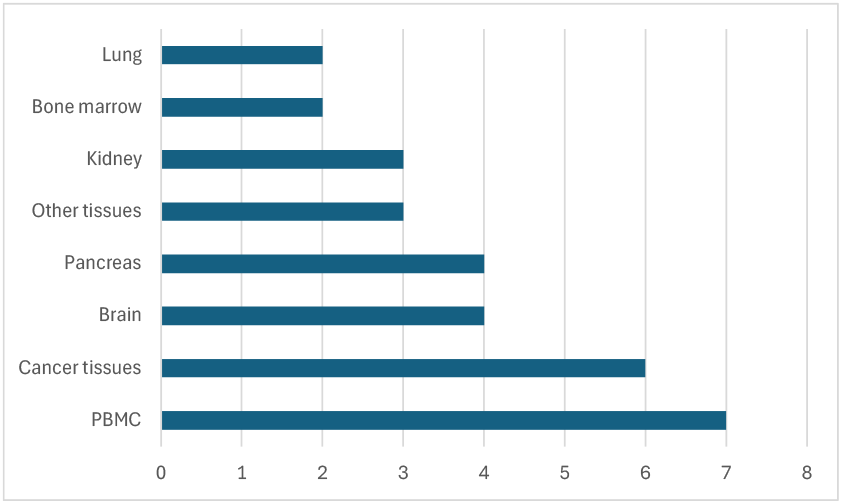
Distribution of Tissue Origins in Deconvolution Datasets

The databases and repositories used by authors to acquire data are also significant. The primary sources are NCBI–GEO^2^, 10x Genomics^3^, ImmPort^4^, and TCGA^5^. NCBI–GEO and 10x Genomics are public repositories for storing high–throughput gene expression data, with the latter specifically focused on scRNA–seq data. ImmPort is an immunology–focused repository, offering access to clinical and experimental data, while TCGA provides genomic, epigenomic, transcriptomics, and clinical data from patients across more than 20 types of cancer.

In Fig. 9, the ‘Others’ category includes databases such as Zenodo^6^, TISCH^7^, Single Cell Portal^8^, Allen Brain Atlas^9^, cBioPortal^10^, GDC Data Portal^11^, LinkedOmics^12^, GTEx^13^, CGGA^14^, EGA^15^and cellxgene^16^. These are grouped together under ‘Others’ as each is mentioned in only one study.

**Fig. 9:**
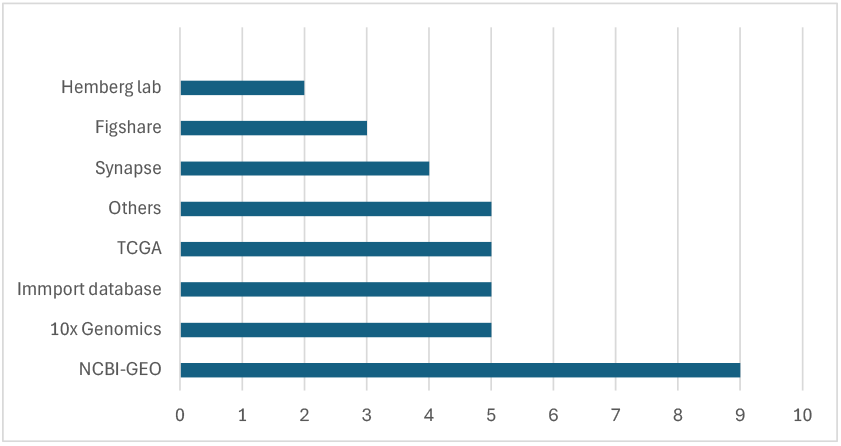
Data Repositories Used in Deconvolution Studies.

After a detailed analysis of each article, it was found that certain datasets are more commonly used across multiple studies. In Table 5 we summarize the most frequently used scRNA–seq datasets and their characteristics, while Table 6 provides the details on each of the bulk RNA–seq datasets.

**Table 5.**
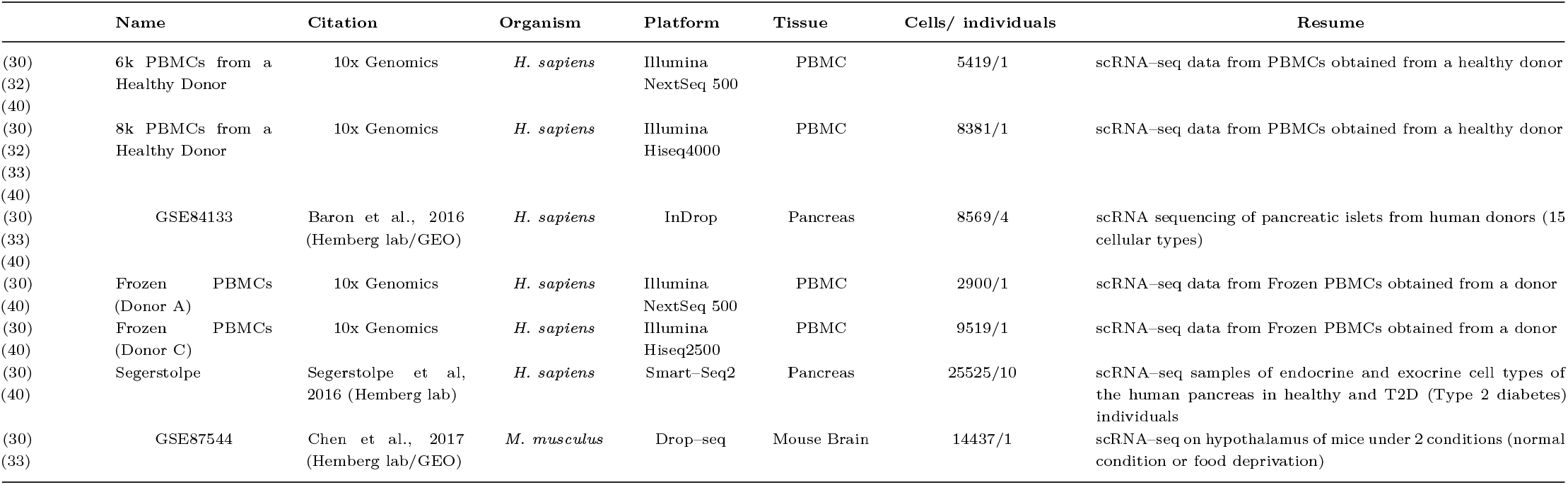
Frequently Used scRNA–seq Datasets and Their Characteristics.

**Table 6.**
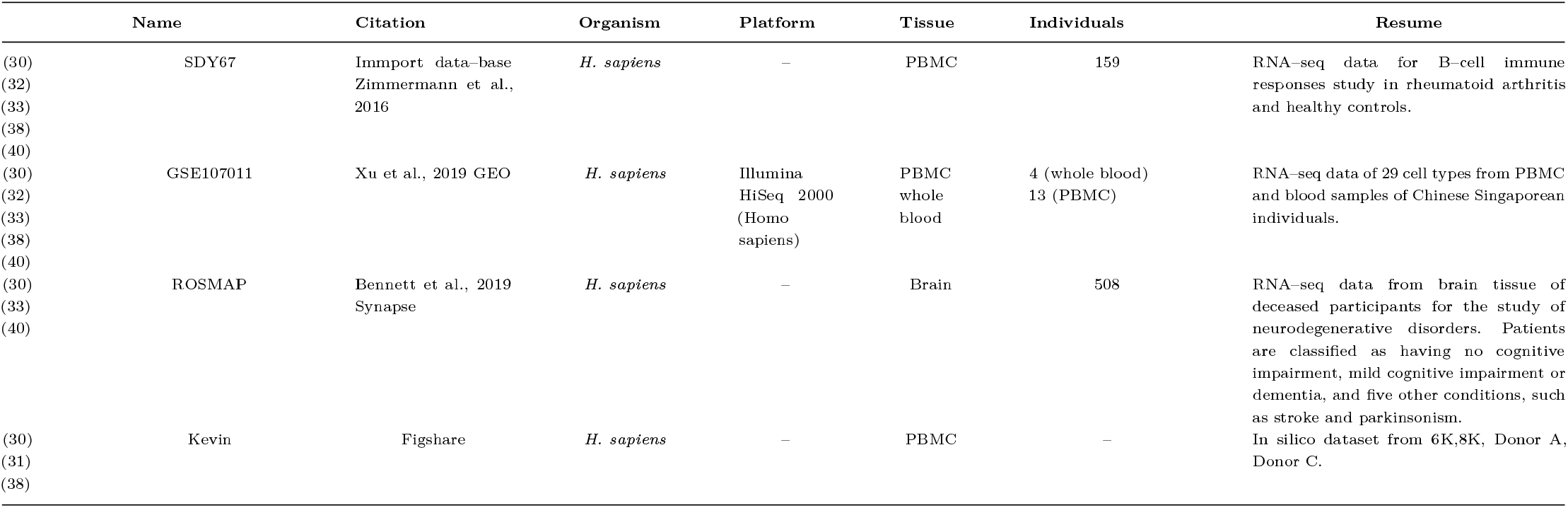
Frequently Used Bulk RNA–seq Datasets in Deconvolution Studies.

### Preprocessing techniques for transcriptomics data

Before deconvoluting transcriptomics data, it is important to consider the preprocessing methods used, as this is a crucial step. In this section, we explore the methodology followed in the preprocessing of scRNA-seq, bulk RNA-seq, and pseudobulk data generation in each of the selected studies we reviewed.

Depending on the specific stage of the workflow within each study, such as constructing pseudobulk data or annotating cell types, the preprocessing of scRNA-seq data can generally be divided into two main stages. The first stage focuses on quality control and normalization. Quality control involves identifying and removing defective data, such as low-quality cells, doublets, and genes with low expression, which could otherwise lead to biased results. Normalization adjusts for systematic and technical variations, such as differences in sequencing depth across cells, enabling accurate comparisons of cells and genes. Effective normalization methods are essential for ensuring that differences observed in gene expression reflect biological variability rather than technical artifacts. After completing these steps, the data are ready for the pseudobulk formation. In Table 3 we outline the procedures employed in each of the studies we reviewed. It is interesting to note that although some studies, such as Scaden, TAPE and DISSECT, work with the same datasets, they present a lack of consistency in the criteria used for quality control. For example, the selected thresholds can significantly differ in terms of the minimum number of genes per cell, the minimum number of cells required for a gene to be considered significant, or the maximum allowable proportion of mitochondrial genes. While the precise effects of these differing criteria remain still unclear, they do influence in the formation of pseudobulk samples and, consequently, in the training of the models.

The second stage is particularly important for cell annotation, which is part of six out of the ten studies that use scRNA–seq data. This processing continues with the calculation of highly variable genes (HVGs), which are then compared to marker genes from the studied cell types. These markers can come from databases like CellMarker^17^for manual annotation or be used for semiautomated annotations through tools like scClassify^18^or CHETAH^19^, which usually require some manual checking afterwards (43). The criteria for selecting HVGs can vary widely, encompassing thresholds for variance and the total number of genes considered. A larger pool of HVGs generally provides more information, leading to more accurate annotations. Furthermore, differences in clustering criteria are evident, particularly with regard to resolution; higher resolution often leads to a greater number of clusters, resulting in the identification of more cell types and a higher level of detail.

Once the scRNA–seq data is processed, the next crucial step is forming pseudobulk samples. This process is vital because DL models require substantial amounts of bulk RNA–seq data. While many real RNA-seq datasets are accessible to the scientific community, they often suffer from limited size or unreliable annotations. Table 7 highlights key considerations in this process, such as the total number of cells (*N*) in each pseudobulk sample, the included cell types (*C*), the method used to calculate cell fractions, the application of data bias techniques (e.g., random gene masking through the addition of Gaussian noise or the creation of sparse samples), and whether cells from different individuals are included in the same pseudobulk sample. It is worth noting that for Digitaldlsorter, not only are multiple pseudobulks created, but individual cellular profiles are also simulated using the ZinbWabe tool^20^, resulting in in silico samples composed of a mixture of real and simulated cells. Additionally, within the GBMPurity tool, pseudobulks are generated dynamically during the training process as the model learns, until it reaches convergence. Once the model has converged, no additional pseudobulk samples are generated.

**Table 7.** Criteria for Pseudobulk Sample Formation in Reviewed Articles. The Bias column indicates whether bias elements are included in the final pseudo–bulk. The Intersample Formation column specifies whether cells from different samples or real and simulated cells were combined in a sample.

|  | *N* (number of cells) | *C* (cell types) | *F<sub>c</sub>* (cell fractions) | Bias | Intersample Formation | Pseudobulk Samples |
| --- | --- | --- | --- | --- | --- | --- |
| (29) | 100 | CRC: 10<br>BC: 10 | Truncated uniform distribution and Dirichlet distribution | No | Yes | 17550 (all datasets) |
| (30) | 500 | PBMC: 5<br>Ascites: 6<br>Pancreas: 7<br>Brain: 7 | Random generation with uniform distribution | Yes | No | 32,000 PBMC<br>14,000 pancreas<br>6000 human brain<br>30,000 mouse brain |
| (32) | 500 | 6 | Dirichlet distribution | No | NA | 16,000 |
| (33) | NS | Limbs: 6<br>Bone marrow: 7<br>Lung: 9<br>PBMC:12<br>Brain: 7<br>Pancreas:5 | Dirichlet distribution with prior knowledge | Yes | No | 5000 |
| (34) | NS | 15 | Random generation with uniform distribution | NS | No | 16000 |
| (36) | 1-10,000 | 1-32 | Random generation | Yes | Yes | 10 M |
| (38) | 500 | 5 | Dirichlet distribution | No | No | 11000 |
| (39) | 3000 | 2 | Uniform distribution | Yes | No | 3000 |
| (40) | 100 | Pancreas: 6<br>PBMC: 18<br>Kidney: 9<br>Brain: 10 | Uniform distribution | No | No | 1, 000XC |
| (41) | 200-400 | NA | Random generation | No | No | 197 |
| (42) | 15 | 4 | Random generation | No | No | 4400 |

Lastly, both real and synthetic RNA-seq data require preprocessing before being input into deconvolution tools. This includes normalizing the count table and selecting the most informative genes, essentially HVGs that will serve as inputs for the model. Like with scRNA–seq data preprocessing, Table 3 shows variations in normalization and gene selection criteria. This is particularly important, as it defines how much information the model will have for deconvolution. There is considerable disparity in practice: for instance, Scaden uses approximately 10,000 genes, while CONVdeconv works with fewer than 2,000 genes.

Additionally, a common practice is to normalize the RNA–seq gene expression data to a range between 0 and 1 using the, so called, min–max scaling, as it has been thoroughly shown that working with normalize datasets enhances the performance of DL models.

It is also important to highlight the software tools used during preprocessing. Out of the 13 papers we have reviewed, seven employed Scanpy^21^(Python), five used Seurat^22^(R), and the remaining studies did not specify the utilized tools.

### Targeted cell types and tissues in deconvolution models

Understanding the specific objectives and use cases for which each deconvolution model is developed is essential for selecting the most appropriate tool for a specific study. These models are typically divided into two main categories: those aimed at addressing specific clinical challenges and those developed to overcome computational constraints. Users choose the tool that best fits their biological or analytical goals, taking into account both their specific objectives and the origin of the data. This distinction provides valuable guidance in making an informed decision.

The first category includes general purpose models such as DeepDecon, aimed at detecting tumor cells; Digitaldlsorter, which performs deconvolution across various cancer types; and Yan et al., 2023 designed a tool for monitoring cancer progression. Additionally, there are more specialized models: (41) focuses on immune system– related changes in mouse Alport syndrome, while GBMPurity was developed to estimate tumor purity in glioblastomas.

### Clinical purposes

In table 3 we show the different types of tissues that are used for training and evaluation, as well as the number of cell types that each model can identify in each kind of tissue. It is noteworthy that most of the deconvolution models are focused on blood tissue analysis, particularly on Peripheral Blood Mononuclear Cells (PBMCs). This focus is not incidental, as PBMCs are of significant and broad interest in biomedical research due to the crucial role they play in the immune system (44). The relative proportion of different cell types in PBMCs can reflect an individual immune status and provide valuable insights in contexts such as early disease detection and immune response monitoring, as well as they are useful for tailoring specific treatments for diseases like cancer or chronic infections. To illustrate the complexity of PBMCs cellular types and the challenge in its classification, we present in fig. 10 a comprehensive overview of PBMCs origin and subtypes.

**Fig. 10:**
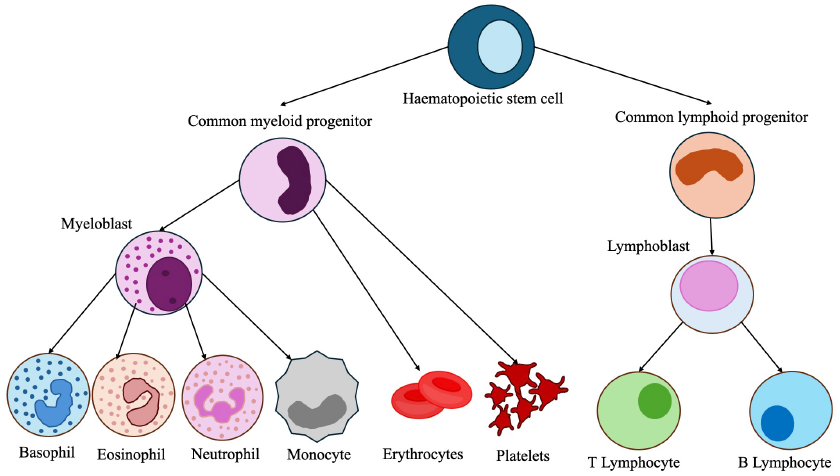
Overview of the different cell types that arise from hematopoietic stem cells through the process of hematopoiesis. The diagram illustrates the differentiation pathways leading to the formation of key blood cell lineages, including myeloid and lymphoid progenitors. The myeloid lineage gives rise to erythrocytes, platelets, monocytes, and granulocytes (basophils, eosinophils, and neutrophils), while the lymphoid lineage differentiates into T and B lymphocytes.

In addition to immune system cell types, some models, such as Digitaldlsorter and DeepDecon, focus exclusively on the study of cancerous samples. The first focuses on the deconvolution of tissues affected by two different types of cancer, offering detailed insights into the tumor state through cellular proportions, while the second aims to estimate the proportions of cancerous cells more generally. These models analyze tumor heterogeneity and its progression, including metastasis.

Other tissues that are examined by several of these models include pancreatic tissue, whose graphical depiction can be seen in Fig. 11, which is critical for analyzing cellular heterogeneity, which is of vital importance in diseases such as diabetes or pancreatic cancer. Furthermore brain tissue has received special attention due to its relevance in research for neurodegenerative diseases such as Alzheimer and other kind of dementias.

**Fig. 11:**
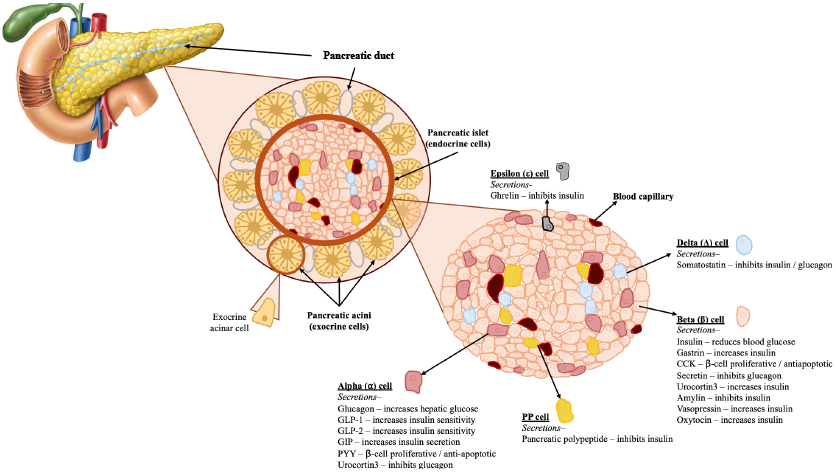
Cellular Heterogeneity in Pancreatic Tissue. The diagram illustrates the structural organization of the pancreas, highlighting key components such as the pancreatic duct, pancreatic islets, and acini. It also depicts the diverse cell types within these structures, emphasizing the complexity of pancreatic tissue architecture.

In the context of pancreas analysis, current deconvolution efforts primarily focus on endocrine cell types, such as alpha, beta, delta, and epsilon cells. While these efforts are valuable for understanding normal pancreatic function, they often overlook the broader cellular landscape of pancreatic tissue, especially relevant in cancerous contexts. A more comprehensive approach should also include other key cell types, such as stromal, immune, and malignant cells, to fully capture the complexity of the tumor micro–environment. These additional cell types play a critical role in disease progression and influence the interactions with endocrine cells. The lack of tools capable of deconvolving the full spectrum of cellular populations limits our ability to fully understand the cellular heterogeneity in pancreatic ductal adenocarcinoma (PDAC), for example.

On the other hand, in the context of clinical applications, the granularity of deconvolution models is of vital importance. Significant differences emerge depending on the tool used, particularly in the resolution of PBMC subclasses. Some tools, such as Autoptcr, can only distinguish two types of T lymphocytes and a single class of B lymphocytes within lymphoblast-derived cells. Others tools, like HASCAD and NNICE, are limited to identifying five PBMC cell types. In contrast, other models such as TAPE can differentiate up to 12 distinct PBMC types, while DISSECT stands out by resolving up to 18 cell types. Higher– resolution methods provide more detailed characterization of cellular subpopulations, facilitating the identification of maturation stages, functional assessments, and a more nuanced understanding of cellular heterogeneity. This enhanced granularity can be critical for clinical decision-making, particularly in monitoring disease progression and therapeutic responses.

### Biological Computational Challenges

The second category includes tools developed to address computational challenges without being tied to a specific biological or clinical context. These solutions encompass novel data generation strategies, such as the pipeline proposed by DAISM-DNN, and the application of innovative DL architectures like TAPE, previously unexplored in this domain. These tools aim to offer adaptable solutions applicable to a wide variety of tissues and cell types, as demonstrated in UCDBase.

The DAISM (Data Augmentation through In Silico Mixing) method tries to enhance the training of deep learning models for estimating cell type proportions from bulk RNA-seq data. To overcome the challenge of limited training data, DAISM generates synthetic samples by in silico mixing of calibration datasets with known cell type fractions. Starting with a small set of real samples, the method creates randomized mixtures of cell types, enabling the development of prediction models that are tailored to specific datasets. This approach helps to avoid cross-platform variability by training exclusively on the augmented dataset.

In HASCAD, the authors propose a versatile model capable of deconvoluting cell types across various transcriptomics data modalities, facilitating the analysis of complex cellular mixtures from different tissues. This approach is particularly valuable for understanding cellular composition and dynamics in diverse biological scenarios. By handling heterogeneous samples, the model offers significant advantages for research into conditions such as idiopathic pulmonary fibrosis, type II diabetes mellitus, and multiple sclerosis.

Table 8 details the cell types by tissue that the tools reviewed here are capable of deconvolving.

**Table 8.**
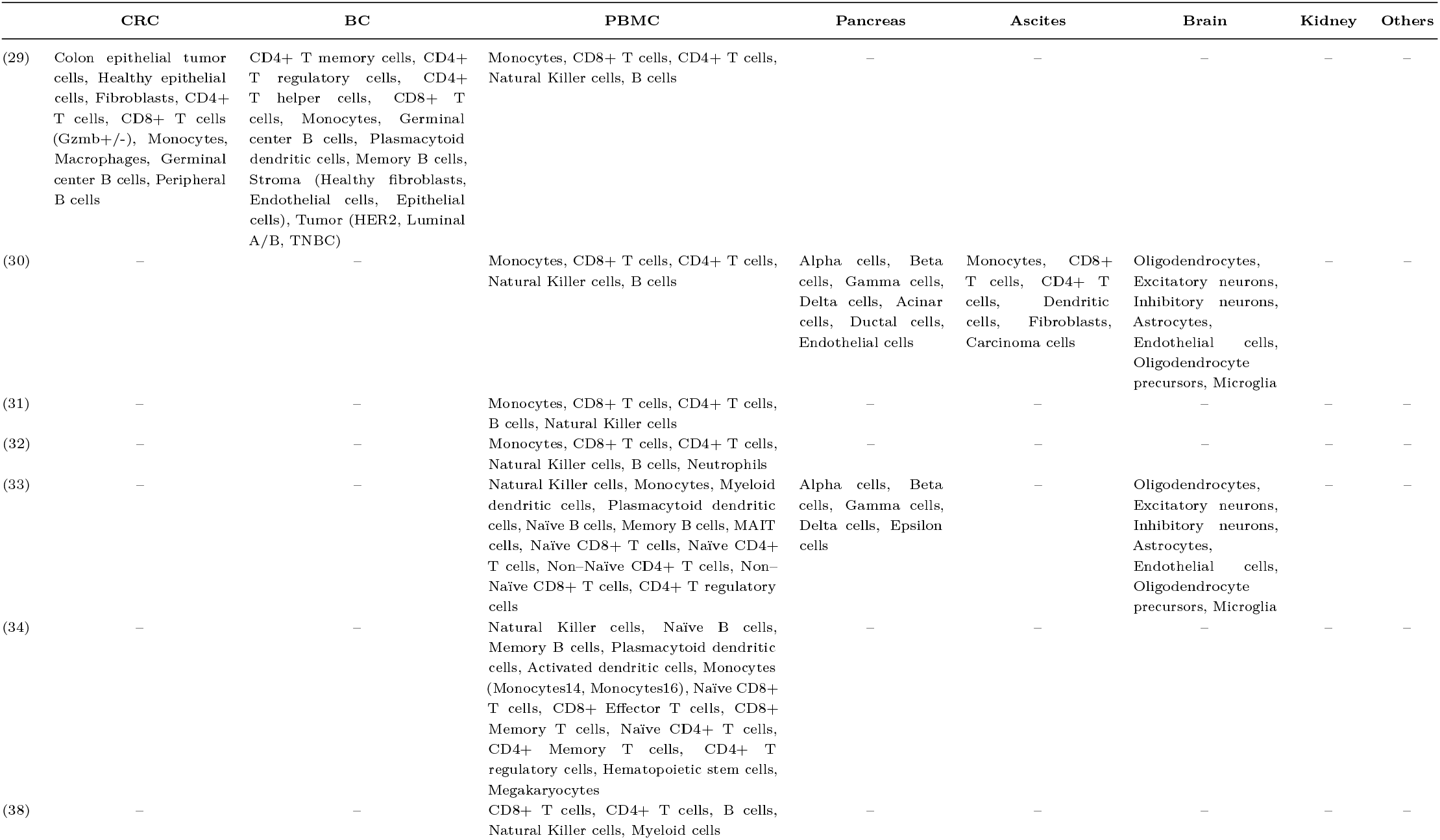
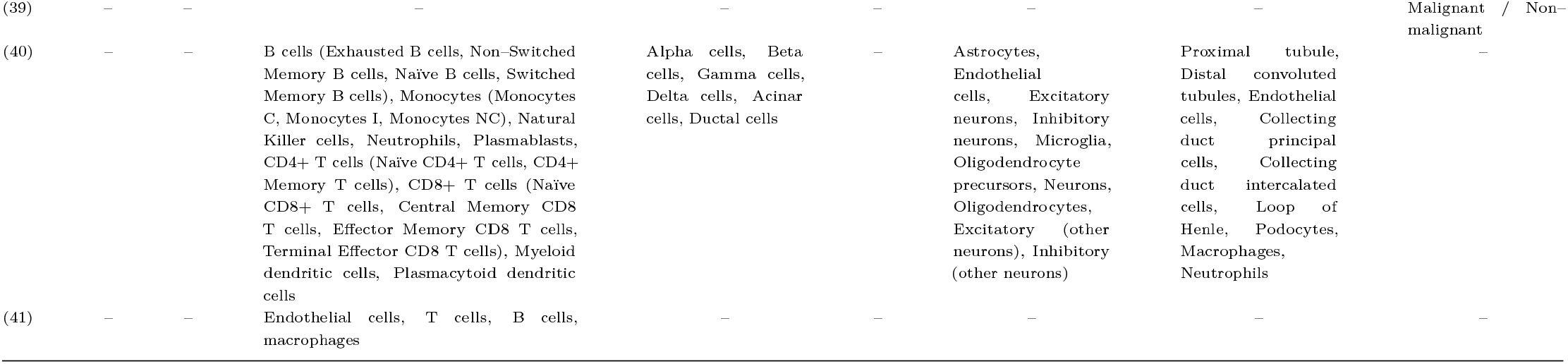
Comparison of deconvoluted cell types across different tissues and tools. The cell types in parentheses are subtypes of the higher-level cell type identified by the tool.

### Predominant DL Architectures: design, training and optimization pipelines

Most DL–based deconvolution tools employ MLP architectures, accounting for 11 out of the 13 tools analyzed (85%). Only three tools utilize different architectures:Autoptcr implements a CNN with four one-dimensional convolutional layers followed by an MLP with a single hidden layer. TAPE, on the other hand, employs a standard autoencoder consisting of two symmetric components (encoder and decoder), each with four fully connected layers; and CONVdeconv features a novel hybrid architecture.

Regarding the development frameworks used, six tools (46%) were developed in TensorFlow, four (31%) in PyTorch, and the remaining three do not specify their development framework.

MLP-based tools employ different strategies. For instance, Scaden and HASCAD use a framework with three distinct MLPs, each estimating cellular proportions. The final prediction is derived from a weighted average of these estimates. In NNICE, it is introduced the use of a neural network model called Deep Quantile Neural Network (DQNN). Unlike traditional neural networks, which predict only the expected value of an output variable, DQNN estimates multiple quantiles of the output variable distribution, which is particularly useful for understanding uncertainty and possible outcome ranges. NNICE calculates cell proportions individually, using one MLP per target cell type for each defined quantile, specifically analyzing 5 quantiles and 6 cell types. DQNN MLPs differ slightly from the classical model by employing a quantile–specific loss function that penalizes deviations asymmetrically according to the desired quantile.

DeepDecon is an iterative DL model to enhance prediction accuracy. It adjusts predictions based on discrepancies between predicted and actual malignant cell fractions, utilizing 55 MLPs, each with four hidden layers, operating across different malignant cell fraction ranges. Lastly, DISSECT incorporates a dual loss function (Tabla 4), combining a reconstruction loss, which measures the model accuracy in reproducing input data, with a consistency loss that encourages similar results between real and simulated RNA–seq samples.

CONVdeconv introduces a hybrid model that combines multiple architectural components, including a convolutional layer, a two– layer MLP, and an attention block, representing a more advanced approach to cellular deconvolution.

The remaining tools rely on simpler MLP architectures, ordered by increasing complexity. Yan et al., 2023 (37) employed a single hidden layer architecture, followed by GMBPurity with two hidden layers. Both DAISM-DNN and Digitaldlsorter feature three hidden layers, while UCDBase incorporates the most complex architecture with four hidden layers.

### Structure of input and output layers

All tools process transcriptomic data from both real bulk RNA-seq samples and in silico simulations as numerical vectors, containing as many values as the number of genes found in the sample. Consequently, the number of neurons in the input layer varies depending on the selected tool and, within each tool, on the tissue being deconvolved, as the number of expressed genes differs across tissues. This particularity may necessitate adjustments to the neural network within the same tool, which can complicate its usability and limit its versatility. For example, in Digitaldlsorter method, the transcriptomics vector for BC samples contains expression data for 34145 genes, whereas for CRC samples, the vector includes 23039 genes. Similarly, the number of neurons in the output layer must be adjusted when the deconvolution target changes, as different cell types will be expressed depending on the tissue. For instance, in TAPE, the model considers 7 cell types for brain tissue, 5 for pancreatic tissue, and 12 for PBMCs.

It is important to note that the amount of input data does not always directly correlate with the granularity achieved in deconvolution. This becomes evident when comparing Scaden and HASCAD, both employing similar architectures with three MLP layers. The former uses data from 10000 genes to deconvolute 5 PBMC cell types, extending to a maximum of 7 types in pancreatic and brain tissues. In contrast, the latter achieves deconvolution of up to 15 PBMC cell types with data from only 2371 genes. Notably, GBMPurity, despite using nearly 6000 genes, provides only a single tumor purity estimate.

### Models training

When it comes to model training, significant differences emerge. Tools such as Digitaldlsorter, Scaden, and TAPE, which are capable of deconvoluting multiple tissue types, determine their final hyperparameters using the grid search technique to optimize performance. For Digitaldlsorter, this process relies solely on data from CRC patients; Scaden uses PBMC and ascites data, while TAPE also works with PBMC data. This targeted hyperparameter tuning approach may restrict model performance when applied to tissues beyond those used for optimization. In contrast, models such as CONVdeconv and GBMPurity focus exclusively on a single tissue type, tailoring hyperparameter selection specifically for that use case.

Furthermore, the availability of training datasets differs across tissues within the same model, leading to inconsistencies in training. For example, methods such as Digitaldlsorter, Scaden, TAPE, and DISSECT operate under different training conditions. The number of training epochs, in particular, differs significantly between approaches: HASCAD is trained for 20 epochs, Digitaldlsorter for 50 epochs, and Scaden for 5000 steps. DISSECT, on the other hand, is trained for 5000*k* epochs, where *k* corresponds to the number of target cell types. Notably, GBMPurity trains the model until convergence, allowing the network to reach an optimized stable state.

In terms of architecture, Table 4 illustrates that most models use the SoftMax activation function in the output layer. This function converts a vector of real numbers into a probability distribution, commonly used in multiclass classification models. It ensures the output probabilities sum to one, with the predicted class corresponding to the highest probability in the output vector. The optimizer used across all models is Adam, with the only variation being the learning rate.

Despite the classification elements in the models, the core task remains regression, as indicated by the employed loss functions. Common regression losses include L1, Mean Squared Error (MSE), Mean Absolute Error (MAE), Root Mean Squared Error (RMSE) and Pearson Correlation Coefficient (r). The only exception is Kullback–Leibler divergence, used by Digitaldlsorter, which measures the difference between two probability distributions and is typically used in classification tasks to assess the divergence between actual and predicted classes.

These considerations are crucial when evaluating the training processes and outcomes of these models. The combination of regression and classification tasks may impact overall performance, a topic that will be discussed further in the next section.

### Benchmarking

One key challenge in evaluating DL models for deconvolution lies in comparing their performance against both traditional and neural methods. Most benchmarked tools are non-neural due to their established use and broader prevalence, with CibersortX (CSx) (45) and MuSiC being the most common, appearing in eight of the thirteen reviewed publications. Less frequently used tools include CIBERSORT (CS), quanTIseq (46), and MCP-Counter (47), found in four and three studies, respectively. In contrast, DL-based tools, though more recent and innovative, remain less numerous. Despite the growing interest in DL models, they are not always benchmarked against other neural-based methods. For instance, Scaden, which is the leading DL method, is only referenced in seven out of the thirteen studies for comparison.

When comparing different deconvolution algorithms, it is important to consider tools requirements. For instance, MCP– Counter relies on specific gene signature matrices, while MuSiC requires scRNA–seq reference data. Neural methods, such as Scaden, need bulk RNA–seq data for network training. Additionally, each tool is designed with specific objectives, which determine the types of tissues they can analyze and their resolution, as discussed in the previous section.

Typically, the first comparisons are conducted using simulated datasets, followed by real datasets. Various metrics are employed to evaluate model accuracy, such as error magnitude (MAE, RMSE, MSE) and correlation (CCC, r) between real and predicted values. Initially, Digitaldlsorter evaluated its performance through correlation plots comparing its results with those from traditional tools such as TIMER, ESTIMATE, EPIC, and MCP–Counter. At that time, DL–based deconvolution tools had not yet emerged, making these comparisons the standard for evaluation. Rather than focusing on the accuracy of the tool, the evaluation focused on how closely its results aligned with those of other established methods in the field.

Methods such as Autoptcr, HASCAD, NNICE, or the one proposed by Yan *et al* (37) do not compare their performance with other deconvolution neural models. Autoptcr paper, for instance, performs its benchmarking against CPM (48), CS, CSx, and MuSiC using four simulated PBMC datasets and one real PBMC dataset, showing the highest correlation and lowest error on simulated data, although all tools exhibited low correlation (< 0.5) when deconvoluting the real dataset. HASCAD tool was evaluated against CSx and quanTIseq using nine real PBMC datasets, outperforming CSx and matching quanTIseq performance while also being able to analyze six additional cell types. Meanwhile, Yan et al., 2023 method surpassed an NNLS–based tool in all simulations across various tissues (brain, breast, colon, kidney, liver, and lung), achieving correlation coefficients (r-values) of 0.98 and 0.79, respectively. Additionally, NNICE approach demonstrated superior correlation metrics compared to TIMER (49), quanTIseq, EPIC, MCP–Counter, and CS using both simulated and real PBMC data, showing correlations of 0.7.

The DAISM-DNN method was compared against several deconvolution tools, including Scaden, MuSiC, CS, CSx, ABIS (50), EPIC, quanTIseq, MCP Counter, and xCell, using the real RNA-seq dataset SDY67. It outperformed all methods, achieving the lowest RMSE and highest correlation (r, CCC) across 30 permutation tests, with particular superiority over Scaden. Further analysis was conducted to determine if DAISM-DNN’s superior performance was solely attributed to the training data generated by the DAISM method. When both Scaden and DAISM-DNN were trained on simulated PBMC data augmented with SDY67 calibration data, their performances became comparable, with improvements observed in both models.

Additionally, TAPE was benchmarked against Scaden, RNASieve (51), CSx, DWLS (52), MuSiC, and Bisque (53), using simulated bulk data from the Tabula Muris Atlas in three scenarios: normal, rare, and similar. Neural methods were more robust in all three scenarios, but DWLS outperformed them in the normal scenario. On real PBMC and brain datasets, TAPE exhibited the highest MAE values and lowest variance, while Scaden had higher correlation values.

In the development of DeepDecon, the benchmarking was again performed with Scaden, CSx, Bisque, ESTIMATE, MuSiC, MEAD (54), RNA–Sieve, and an NNLS–based model, using datasets from AML^23^, HNSCC^24^, and neuroblastoma patients. DeepDecon outperformed its counterparts in 11 of 15 AML datasets and achieved the lowest RMSE and highest correlations for HNSCC and neuroblastoma datasets.

The UCDBase evaluation of RNA–seq deconvolution included SCDC (55), MuSiC, and Scaden. The results showed that the method achieved a mean r–value score of 0.68 when deconvolving 96 cell mixtures.

In CONVdeconv, comparisons are made using simulated data with two DL–based deconvolution tools: Scaden and DestVI (56), a spatial transcriptomics deconvolution model. Notably, both Scaden and DestVI exhibit negative *r*–values, whereas CONVdeconv achieves an *r*–value of 0.86. Additionally, CONVdeconv attained the lowest RMSE and MAE values (RMSE = 0.07, MAE = 0.06). GBMPurity benchmarking is performed against MuSiC, CIBERSORTx, PUREE (57), and Scaden. A comparison across two bulk RNA–seq datasets shows that the PUREE (MAE = 0.102, RMSE = 0.123, r = 0.803, CCC = 0.701) and GBMPurity (MAE = 0.128, RMSE = 0.160, r = 0.757, CCC = 0.743) models yield the best metrics when estimating tumor purity in glioblastoma within the EORTC and TCGA datasets, respectively. Scaden performs the worst in both cases.

Finally, DISSECT expanded neural tool comparisons to include Scaden, TAPE, and an MLP, as well as traditional methods such as MuSiC, CSx, BayesPrism (54), and b–MIND (58). Metrics such as JSD (Jensen-Shannon Divergence) were used for additional evaluations, and DISSECT consistently emerged as the top performer across PBMC, brain, pancreas, and kidney datasets, though the second–best tool varied by tissue type: TAPE for kidney, the MLP for brain, and Scaden for PBMC.

In summary, neural-based tools are increasingly featured in comparative deconvolution studies, highlighting their growing importance in the field. In this benchmarking evaluations, DL models generally outperform traditional methods, although their effectiveness varies across tissue types and datasets, emphasizing the need for context-specific assessments.

## Discussion

Deep learning-based deconvolution methods have demonstrated significant potential in bulk RNA-seq analysis. However, our review highlights several fundamental challenges that remain unresolved, particularly concerning model design, data preprocessing, and generalization across different tissues. Addressing these limitations is crucial for ensuring the reliable application of these methods in clinical and biomedical research.

### Computational Field

One of the most relevant issues arising from our review is the existing controversy surrounding the categorization of the cellular deconvolution problem as either a regression or classification problem. While most of the Deep Learning models employ loss functions typical of regression problems, such as mean squared error (MSE), a considerable number of implementations use the SoftMax function in the output layer instead. This latter function turns predictions into probability distributions over different classes, creating a methodological duality. On one hand, the primary goal is to predict cellular proportions, aligning with a regression approach; on the other hand, the use of SoftMax introduces a probabilistic interpretation similar to a classification problem. Investigating the impact of this choice on the accuracy of Deep Learning models is crucial, as well as the consideration of whether the deconvolution problem should be strictly addressed as regression task or if a reformulation as a classification one might be more suitable in certain contexts.

Another critical aspect is whether the application of a given neural network architecture, trained with data from a specific cell type, to deconvolve different tissues without making modifications to the internal structure of the model is a well suited problem. In this regard, tools like Scaden train their model using data from PBMCs and apply the resulting model to deconvolve pancreatic or brain tissues. This strategy raises questions about the validity of assuming that gene expression patterns are comparable across these different contexts, given that gene expression and regulatory cell interactions are highly dependent on the considered tissue. This tissue–dependency suggests that the direct transfer of a model trained within an specific tissue context to analyze the composition of different tissues may not be appropriate without a more thorough adjustment. We would like to emphasize the necessity of a robust study on the adaption of each model to the specific characteristics of each tissue.

Furthermore, it is noteworthy that even within the same tool, the number of samples used to train the model varies significantly between different tissues. This potential inconsistency could affect the robustness and generalization of the models, suggesting that greater standardization in data usage and training methodology is crucial to enhance the effectiveness of deconvolution approaches.

An additional significant challenge arises when in we want to compare the effectiveness between different deconvolution models. Each of the revised tools handles data differently and presents varying levels of accuracy, leading then to discrepancies in both their inputs (genes) and outputs (cellular proportions or classes). Furthermore, the existence of diverse validation methods complicates this comparison, as there is no single standard criterion for evaluating and comparing the results obtained by each model. This lack of homogeneity sticks out the need of establishing uniform criteria and protocols in the field of cellular deconvolution.

Finally, although most current models are based on the same architecture as Scaden, an MLP network, recent efforts are being made to implement more sophisticated models, such as Autoptcr or TAPE, which are based on CNNs and Autoencoders, respectively. This evolution indicates that the field is still in its early stages of development, leaving room for innovation. This rather straightforward use of simple models leave a lot of room for improvement, so future work in this discipline may include the implementation of more complex architectures, based on attention mechanisms or graph networks, which may be more suited to encode the rich variability inherent to transcriptomics data. Another rather weak point for deconvolution tools training is the scarcity of training data. A great amount of effort has been devoted lately to build reliable generators of synthetic scRNA–seq data, so more realistic pseudobulks can be implemented to make more robust tools (59), (60), (61).

### Biological Field

In the biological realm, as we already mentioned above, the scarcity of RNA–seq data represents a significant obstacle for constructing robust models. Currently, there is a reliance on generating in silico (pseudobulk) data that attempts to simulate RNA–seq data using scRNA–seq data. However, intrinsic differences between these data types, which are not yet fully understood, could affect the proper training of the models. Additionally, the process of generating these pseudobulks varies considerably between studies, as we have noted in thgis review, sticking out the lack of standardization in this aspect.

Moreover, access to annotated scRNA–seq data, necessary for generating training pseudobulks, presents an additional challenge. Often, these data are unlabeled, requiring a labor–intensive, complex process that may be difficult to reproduce without the original pipeline. Therefore, it is essential to develop a consensus on best practices for the generation and preprocessing of transcriptomics data, as this could improve the consistency and comparability of results obtained with Deep Learning models.

Regarding the target tissues in the analyzed studies, it is notable that there is a predominance of samples from PBMC, with only one article addressing deconvolution in two distinct types of cancer. This trend limits the scope of deconvolution tools and raises questions about their applicability and effectiveness in broader clinical contexts.

Future directions in this field include the need to achieve a representation of RNA–seq data that provides greater insight into the dynamics and interdependence of cells. Cellular states play a crucial role in the variability of gene expression, and the emergence of one cell type can influence the expression of other types. Understanding these interconnections is essential to enhance the accuracy of deconvolution models.

Furthermore, it is crucial to develop models capable of deconvolving a wide variety of tissues, rather than being limited solely to PBMC samples, as has been the predominant approach thus far. Expanding the spectrum of tissues addressed could have a significant impact on clinical practice, facilitating the study of various diseases and allowing for a more comprehensive analysis of how cellular proportions evolve in patient samples over time.

Lastly, it is important to emphasize the need for standardization in data acquisition for training and validation methods, which could clarify the utility of deconvolution models in biomedical research.

## Conclusions

In this review, we have examined several different Deep Learning–based cell deconvolution tools, highlighting their key methodological and conceptual challenges, and the potential future research directions in this rapidly evolving field.

A major common issue we have encountered through this revision is the wide variability in the outputs generated by each tool, including not only the differences in the selected cell types that are identified but also in the format of the presented results. This heterogeneity rather complicates any direct comparison between different methods, therefore emphasizing the necessity of standardized evaluation criteria to enable an objective benchmarking.

There is an ongoing debate concerning the best approach to tool development. While some studies aim to create generalized models that can adapt to samples coming from different tissues and obtained in diverse experimental conditions, others studies sustain that focusing on obtaining specialized models optimized for specific applications is better suited to the variable nature of transcriptomics data. Generalized models offer the advantage of broad applicability, as they can be used across multiple datasets and conditions without requiring major architecture changes. This flexibility is particularly useful in large–scale studies where samples come from different sources, as well as in clinical settings where robustness across various patient–derived samples is crucial. However, this generalizability often comes at the cost of decreased precision, as these models may not fully capture tissue-specific transcriptomics signatures.

On the other hand, specialized models tailored to specific tissues or experimental settings can achieve higher accuracy by leveraging domain-specific features and optimizing hyperparameters accordingly. These models are particularly beneficial in applications where fine–grained cellular resolution is needed, such as tumor microenvironment studies or immune profiling in disease contexts. Nevertheless, their applicability is more limited, as they may require retraining or fine-tuning when applied to different datasets, reducing their usability in diverse research scenarios.

Both strategies present different compromises, and their selection depends on the intended application, the availability of training data, and the desired balance between accuracy and generalizability. Future developments in this field may involve hybrid approaches that integrate the strengths of both methodologies, potentially leveraging meta–learning or transfer learning techniques to enhance adaptability without sacrificing precision.

Moreover, almost all of the developed deconvolution models rely on relatively simple Deep Learning architectures. Henceforth there is still significant room for improvement through the implementation of more sophisticated models capable of better capturing the complexity of transcriptomics data. Additionally, transcriptomics data has traditionally been represented as vectors, yet alternative representations –such as image–based formats or structured data models—-could enhance the predictive power of these alternative models and provide novel insights that can potentially be more accurate in the deconvolution tasks.

Finally, the generation of synthetic data tailored to the specific challenges of cell deconvolution emerges as a potential key strategy for improving model training and validation. The development of well–characterized datasets that accurately reflect the biological heterogeneity of tissues would enhance model robustness and broaden their applicability in both research and clinical settings.

## Supporting information

Latex files

## Author contributions statement

A.L.R. has performed the PRISMA methodology to search for the significative studies to include in the review, and written the manuscript, and created the figures and comparative analysis.

J.M.S.V has supervised the elaboration of the manuscript and have contributed to the final version. V.J.S-A.L has supervised the elaboration of the manuscript and have contributed to the final version.

## Acknowledgments

The authors would like to warmly thank Alberto Nogales for our enlightening discussions and his invaluable suggestions along the way. They also want to express their gratitude to Maite Iglesias Badiola, Cruz Santos Tejedor, Alberto Lpéz Rosado, and Jose Luis Rodríguez Peralto for their unwavering support to the group.

1 http://www.prisma-statement.org

2 https://www.ncbi.nlm.nih.gov/geo/

3 https://www.10xgenomics.com/

4 https://immport.org/

5 https://www.cancer.gov/ccg/research/genome-sequencing/tcga

6 https://zenodo.org/

7 http://tisch.comp-genomics.org/home/

8 https://singlecell.broadinstitute.org/single_cell

9 https://atlas.brain-map.org/

10 https://www.cbioportal.org/

11 https://portal.gdc.cancer.gov/

12 http://linkedomics.org/

13 https://www.gtexportal.org/home/

14 https://www.cgga.org.cn/

15 https://ega-archive.org/

16 https://cellxgene.cziscience.com/datasets

17 http://xteam.xbio.top/CellMarker/

18 https://github.com/SydneyBioX/scClassify

19 https://github.com/jdekanter/CHETAH

20 https://bioconductor.org/packages/zinbwave/

21 https://scanpy.readthedocs.io/

22 https://satijalab.org/seurat/

23 Acute Myeloid Leukemia

24 Head and Neck Squamous Cell Carcinoma

