## Supplementary material for "Deep Learning based deconvolution methods: a systematic review": Latex files: reference.bib

@article{wang2020,
    author = {Wang, Shiyuan and Zhang, Qi and Yu, Chunlu and others},
    title = {{Immune cell infiltration-based signature for prognosis and immunogenomic analysis in breast cancer}},
    journal = {Briefings in Bioinformatics},
    volume = {22},
    number = {2},
    pages = {2020-2031},
    year = {2020},
    month = {03},
    abstract = {Breast cancer is one of the most human malignant diseases and the leading cause of cancer-related death in the world. However, the prognostic and therapeutic benefits of breast cancer patients cannot be predicted accurately by the current stratifying system. In this study, an immune-related prognostic score was established in 22 breast cancer cohorts with a total of 6415 samples. An extensive immunogenomic analysis was conducted to explore the relationships between immune score, prognostic significance, infiltrating immune cells, cancer genotypes and potential immune escape mechanisms. Our analysis revealed that this immune score was a promising biomarker for estimating overall survival in breast cancer. This immune score was associated with important immunophenotypic factors, such as immune escape and mutation load. Further analysis revealed that patients with high immune scores exhibited therapeutic benefits from chemotherapy and immunotherapy. Based on these results, we can conclude that this immune score may be a useful tool for overall survival prediction and treatment guidance for patients with breast cancer.},
    issn = {1477-4054},
    doi = {10.1093/bib/bbaa026},
    url = {https://doi.org/10.1093/bib/bbaa026},
    eprint = {https://academic.oup.com/bib/article-pdf/22/2/2020/36670086/bbaa026.pdf},
}

@ARTICLE{zhu2022,
AUTHOR={Zhu, Zheng  and Li, Guoyin  and Li, Zhenning and others},
TITLE={{Core immune cell infiltration signatures identify molecular subtypes and promote precise checkpoint immunotherapy in cutaneous melanoma}},
JOURNAL={Frontiers in Immunology},
VOLUME={13},
YEAR={2022},
URL={https://www.frontiersin.org/journals/immunology/articles/10.3389/fimmu.2022.914612},
DOI={10.3389/fimmu.2022.914612},
ISSN={1664-3224},
ABSTRACT={<p>Yutao Wang, China Medical University, ChinaThe tumor microenvironment (TME) has been shown to impact the prognosis of tumors in patients including cutaneous melanoma (CM); however, not all components of TME are important. Given the aforementioned situation, the functional immune cell contents correlated with CM patient prognosis are needed to optimize present predictive models and reflect the overall situation of TME. We developed a novel risk score named core tumor-infiltrating immune cell score (cTICscore), which showed certain advantages over existing biomarkers or TME-related signatures in predicting the prognosis of CM patients. Furthermore, we explored a new gene signature named cTILscore−related module gene score (cTMGs), based on four identified TME-associated genes (<italic>GCH1</italic>, <italic>GZMA</italic>, <italic>PSMB8</italic>, and <italic>PLAAT4</italic>) showing a close correlation with the cTICscore, which was generated by weighted gene co-expression network analysis and least absolute shrinkage and selection operator analysis to facilitate clinical application. Patients with low cTMGs had significantly better overall survival (OS, <italic>P</italic> = 0.002,&lt; 0.001, = 0.002, and = 0.03, respectively) in the training and validating CM datasets. In addition, the area under the curve values used to predict the immune response in four CM cohorts were 0.723, 0.723, 0.754, and 0.792, respectively, and that in one gastric cohort was 0.764. Therefore, the four-gene signature, based on cTICscore, might improve prognostic information, serving as a predictive tool for CM patients receiving immunotherapy.cutaneous melanoma, tumor microenvironment, prognosis, immunotherapy, cTICscore</p>}}

@article{nguyen2024,
    author = {Nguyen, Hung and Nguyen, Ha and Tran, Duc and others},
    title = {{Fourteen years of cellular deconvolution: methodology, applications, technical evaluation and outstanding challenges}},
    journal = {Nucleic Acids Research},
    volume = {52},
    number = {9},
    pages = {4761-4783},
    year = {2024},
    month = {04},
    abstract = {Single-cell RNA sequencing (scRNA-Seq) is a recent technology that allows for the measurement of the expression of all genes in each individual cell contained in a sample. Information at the single-cell level has been shown to be extremely useful in many areas. However, performing single-cell experiments is expensive. Although cellular deconvolution cannot provide the same comprehensive information as single-cell experiments, it can extract cell-type information from bulk RNA data, and therefore it allows researchers to conduct studies at cell-type resolution from existing bulk datasets. For these reasons, a great effort has been made to develop such methods for cellular deconvolution. The large number of methods available, the requirement of coding skills, inadequate documentation, and lack of performance assessment all make it extremely difficult for life scientists to choose a suitable method for their experiment. This paper aims to fill this gap by providing a comprehensive review of 53 deconvolution methods regarding their methodology, applications, performance, and outstanding challenges. More importantly, the article presents a benchmarking of all these 53 methods using 283 cell types from 30 tissues of 63 individuals. We also provide an R package named DeconBenchmark that allows readers to execute and benchmark the reviewed methods (https://github.com/tinnlab/DeconBenchmark).},
    issn = {0305-1048},
    doi = {10.1093/nar/gkae267},
    url = {https://doi.org/10.1093/nar/gkae267},
    eprint = {https://academic.oup.com/nar/article-pdf/52/9/4761/57807901/gkae267.pdf},
}


@article{lecun2015,
	abstract = {Deep learning allows computational models that are composed of multiple processing layers to learn representations of data with multiple levels of abstraction. These methods have dramatically improved the state-of-the-art in speech recognition, visual object recognition, object detection and many other domains such as drug discovery and genomics. Deep learning discovers intricate structure in large data sets by using the backpropagation algorithm to indicate how a machine should change its internal parameters that are used to compute the representation in each layer from the representation in the previous layer. Deep convolutional nets have brought about breakthroughs in processing images, video, speech and audio, whereas recurrent nets have shone light on sequential data such as text and speech.},
	author = {LeCun, Yann and Bengio, Yoshua and Hinton, Geoffrey},
	date = {2015/05/01},
	date-added = {2025-02-03 17:55:46 +0100},
	date-modified = {2025-02-03 17:55:46 +0100},
	doi = {10.1038/nature14539},
	id = {LeCun2015},
	isbn = {1476-4687},
	journal = {Nature},
	number = {7553},
	pages = {436--444},
	title = {{Deep learning}},
	url = {https://doi.org/10.1038/nature14539},
	volume = {521},
	year = {2015},
	bdsk-url-1 = {https://doi.org/10.1038/nature14539}}


@article{jumper2021,
	abstract = {Proteins are essential to life, and understanding their structure can facilitate a mechanistic understanding of their function. Through an enormous experimental effort1--4, the structures of around 100,000 unique proteins have been determined5, but this represents a small fraction of the billions of known protein sequences6,7. Structural coverage is bottlenecked by the months to years of painstaking effort required to determine a single protein structure. Accurate computational approaches are needed to address this gap and to enable large-scale structural bioinformatics. Predicting the three-dimensional structure that a protein will adopt based solely on its amino acid sequence---the structure prediction component of the `protein folding problem'8---has been an important open research problem for more than 50 years9. Despite recent progress10--14, existing methods fall far short of atomic accuracy, especially when no homologous structure is available. Here we provide the first computational method that can regularly predict protein structures with atomic accuracy even in cases in which no similar structure is known. We validated an entirely redesigned version of our neural network-based model, AlphaFold, in the challenging 14th Critical Assessment of protein Structure Prediction (CASP14)15, demonstrating accuracy competitive with experimental structures in a majority of cases and greatly outperforming other methods. Underpinning the latest version of AlphaFold is a novel machine learning approach that incorporates physical and biological knowledge about protein structure, leveraging multi-sequence alignments, into the design of the deep learning algorithm.},
	author = {Jumper, John and Evans, Richard and Pritzel, Alexander and others},
	date = {2021/08/01},
	date-added = {2025-02-03 18:09:19 +0100},
	date-modified = {2025-02-03 18:09:19 +0100},
	doi = {10.1038/s41586-021-03819-2},
	id = {Jumper2021},
	isbn = {1476-4687},
	journal = {Nature},
	number = {7873},
	pages = {583--589},
	title = {{Highly accurate protein structure prediction with AlphaFold}},
	url = {https://doi.org/10.1038/s41586-021-03819-2},
	volume = {596},
	year = {2021},
	bdsk-url-1 = {https://doi.org/10.1038/s41586-021-03819-2}}


@article{carpenter2006,
	abstract = {Biologists can now prepare and image thousands of samples per day using automation, enabling chemical screens and functional genomics (for example, using RNA interference). Here we describe the first free, open-source system designed for flexible, high-throughput cell image analysis, CellProfiler. CellProfiler can address a variety of biological questions quantitatively, including standard assays (for example, cell count, size, per-cell protein levels) and complex morphological assays (for example, cell/organelle shape or subcellular patterns of DNA or protein staining).},
	author = {Carpenter, Anne E. and Jones, Thouis R. and Lamprecht, Michael R. and others},
	date = {2006/10/31},
	date-added = {2025-02-03 18:10:47 +0100},
	date-modified = {2025-02-03 18:10:47 +0100},
	doi = {10.1186/gb-2006-7-10-r100},
	id = {Carpenter2006},
	isbn = {1474-760X},
	journal = {Genome Biology},
	number = {10},
	pages = {R100},
	title = {{CellProfiler: image analysis software for identifying and quantifying cell phenotypes}},
	url = {https://doi.org/10.1186/gb-2006-7-10-r100},
	volume = {7},
	year = {2006},
	bdsk-url-1 = {https://doi.org/10.1186/gb-2006-7-10-r100}}


@article{poplin2018,
	abstract = {DeepVariant uses convolutional neural networks to improve the accuracy of variant calling.},
	author = {Poplin, Ryan and Chang, Pi-Chuan and Alexander, David and others},
	date = {2018/11/01},
	date-added = {2025-02-03 18:11:55 +0100},
	date-modified = {2025-02-03 18:11:55 +0100},
	doi = {10.1038/nbt.4235},
	id = {Poplin2018},
	isbn = {1546-1696},
	journal = {Nature Biotechnology},
	number = {10},
	pages = {983--987},
	title = {{A universal SNP and small-indel variant caller using deep neural networks}},
	url = {https://doi.org/10.1038/nbt.4235},
	volume = {36},
	year = {2018},
	bdsk-url-1 = {https://doi.org/10.1038/nbt.4235}}


@article{newman2015,
	abstract = {A computational method to identify cell types within a complex tissue, based on analysis of gene expression profiles, is described in this paper.},
	author = {Newman, Aaron M and Liu, Chih Long and Green, Michael R and others},
	date = {2015/05/01},
	date-added = {2025-02-04 11:13:48 +0100},
	date-modified = {2025-02-04 11:13:48 +0100},
	doi = {10.1038/nmeth.3337},
	id = {Newman2015},
	isbn = {1548-7105},
	journal = {Nature Methods},
	number = {5},
	pages = {453--457},
	title = {{Robust enumeration of cell subsets from tissue expression profiles}},
	url = {https://doi.org/10.1038/nmeth.3337},
	volume = {12},
	year = {2015},
	bdsk-url-1 = {https://doi.org/10.1038/nmeth.3337}}


@article{yoshihara2013,
	abstract = {Infiltrating stromal and immune cells form the major fraction of normal cells in tumour tissue and not only perturb the tumour signal in molecular studies but also have an important role in cancer biology. Here we describe `Estimation of STromal and Immune cells in MAlignant Tumours using Expression data'(ESTIMATE)---a method that uses gene expression signatures to infer the fraction of stromal and immune cells in tumour samples. ESTIMATE scores correlate with DNA copy number-based tumour purity across samples from 11 different tumour types, profiled on Agilent, Affymetrix platforms or based on RNA sequencing and available through The Cancer Genome Atlas. The prediction accuracy is further corroborated using 3,809 transcriptional profiles available elsewhere in the public domain. The ESTIMATE method allows consideration of tumour-associated normal cells in genomic and transcriptomic studies. An R-library is available on https://sourceforge.net/projects/estimateproject/.},
	author = {Yoshihara, Kosuke and Shahmoradgoli, Maria and Mart{\'\i}nez, Emmanuel and others},
	date = {2013/10/11},
	date-added = {2025-02-04 11:14:35 +0100},
	date-modified = {2025-02-04 11:14:35 +0100},
	doi = {10.1038/ncomms3612},
	id = {Yoshihara2013},
	isbn = {2041-1723},
	journal = {Nature Communications},
	number = {1},
	pages = {2612},
	title = {{Inferring tumour purity and stromal and immune cell admixture from expression data}},
	url = {https://doi.org/10.1038/ncomms3612},
	volume = {4},
	year = {2013},
	bdsk-url-1 = {https://doi.org/10.1038/ncomms3612}}


@article{wang2019,
	abstract = {Knowledge of cell type composition in disease relevant tissues is an important step towards the identification of cellular targets of disease. We present MuSiC, a method that utilizes cell-type specific gene expression from single-cell RNA sequencing (RNA-seq) data to characterize cell type compositions from bulk RNA-seq data in complex tissues. By appropriate weighting of genes showing cross-subject and cross-cell consistency, MuSiC enables the transfer of cell type-specific gene expression information from one dataset to another. When applied to pancreatic islet and whole kidney expression data in human, mouse, and rats, MuSiC outperformed existing methods, especially for tissues with closely related cell types. MuSiC enables the characterization of cellular heterogeneity of complex tissues for understanding of disease mechanisms. As bulk tissue data are more easily accessible than single-cell RNA-seq, MuSiC allows the utilization of the vast amounts of disease relevant bulk tissue RNA-seq data for elucidating cell type contributions in disease.},
	author = {Wang, Xuran and Park, Jihwan and Susztak, Katalin and others},
	date = {2019/01/22},
	date-added = {2025-02-04 11:15:26 +0100},
	date-modified = {2025-02-04 11:15:26 +0100},
	doi = {10.1038/s41467-018-08023-x},
	id = {Wang2019},
	isbn = {2041-1723},
	journal = {Nature Communications},
	number = {1},
	pages = {380},
	title = {{Bulk tissue cell type deconvolution with multi-subject single-cell expression reference}},
	url = {https://doi.org/10.1038/s41467-018-08023-x},
	volume = {10},
	year = {2019},
	bdsk-url-1 = {https://doi.org/10.1038/s41467-018-08023-x}}


@article{wang2014,
    author = {Wang, Niya and Gong, Ting and Clarke, Robert and others},
    title = {{UNDO: a Bioconductor R package for unsupervised deconvolution of mixed gene expressions in tumor samples}},
    journal = {Bioinformatics},
    volume = {31},
    number = {1},
    pages = {137-139},
    year = {2014},
    month = {09},
    abstract = {Summary: We develop a novel unsupervised deconvolution method, within a well-grounded mathematical framework, to dissect mixed gene expressions in heterogeneous tumor samples. We implement an R package, UNsupervised DecOnvolution (UNDO), that can be used to automatically detect cell-specific marker genes (MGs) located on the scatter radii of mixed gene expressions, estimate cellular proportions in each sample and deconvolute mixed expressions into cell-specific expression profiles. We demonstrate the performance of UNDO over a wide range of tumor–stroma mixing proportions, validate UNDO on various biologically mixed benchmark gene expression datasets and further estimate tumor purity in TCGA/CPTAC datasets. The highly accurate deconvolution results obtained suggest not only the existence of cell-specific MGs but also UNDO’s ability to detect them blindly and correctly. Although the principal application here involves microarray gene expressions, our methodology can be readily applied to other types of quantitative molecular profiling data. Availability and implementation: UNDO is available at http://bioconductor.org/packages . Contact:   information:  Supplementary data are available at Bioinformatics online. },
    issn = {1367-4803},
    doi = {10.1093/bioinformatics/btu607},
    url = {https://doi.org/10.1093/bioinformatics/btu607},
    eprint = {https://academic.oup.com/bioinformatics/article-pdf/31/1/137/49011292/bioinformatics\_31\_1\_137.pdf},
}

@article{kang2019,
    doi = {10.1371/journal.pcbi.1007510},
    author = {Kang, Kai AND Meng, Qian AND Shats, Igor AND others},
    journal = {PLOS Computational Biology},
    publisher = {Public Library of Science},
    title = {{CDSeq: A novel complete deconvolution method for dissecting heterogeneous samples using gene expression data}},
    year = {2019},
    month = {12},
    volume = {15},
    url = {https://doi.org/10.1371/journal.pcbi.1007510},
    pages = {1-18},
    abstract = {Quantifying cell-type proportions and their corresponding gene expression profiles in tissue samples would enhance understanding of the contributions of individual cell types to the physiological states of the tissue. Current approaches that address tissue heterogeneity have drawbacks. Experimental techniques, such as fluorescence-activated cell sorting, and single cell RNA sequencing are expensive. Computational approaches that use expression data from heterogeneous samples are promising, but most of the current methods estimate either cell-type proportions or cell-type-specific expression profiles by requiring the other as input. Although such partial deconvolution methods have been successfully applied to tumor samples, the additional input required may be unavailable. We introduce a novel complete deconvolution method, CDSeq, that uses only RNA-Seq data from bulk tissue samples to simultaneously estimate both cell-type proportions and cell-type-specific expression profiles. Using several synthetic and real experimental datasets with known cell-type composition and cell-type-specific expression profiles, we compared CDSeq’s complete deconvolution performance with seven other established deconvolution methods. Complete deconvolution using CDSeq represents a substantial technical advance over partial deconvolution approaches and will be useful for studying cell mixtures in tissue samples. CDSeq is available at GitHub repository (MATLAB and Octave code): https://github.com/kkang7/CDSeq.},
    number = {12},

}


@article{li2019,
	abstract = {In the analysis of high-throughput data from complex samples, cell composition is an important factor that needs to be accounted for. Except for a limited number of tissues with known pure cell type profiles, a majority of genomics and epigenetics data relies on the ``reference-free deconvolution''methods to estimate cell composition. We develop a novel computational method to improve reference-free deconvolution, which iteratively searches for cell type-specific features and performs composition estimation. Simulation studies and applications to six real datasets including both DNA methylation and gene expression data demonstrate favorable performance of the proposed method. TOAST is available at https://bioconductor.org/packages/TOAST.},
	author = {Li, Ziyi and Wu, Hao},
	date = {2019/09/04},
	date-added = {2025-02-04 11:23:55 +0100},
	date-modified = {2025-02-04 11:23:55 +0100},
	doi = {10.1186/s13059-019-1778-0},
	id = {Li2019},
	isbn = {1474-760X},
	journal = {Genome Biology},
	number = {1},
	pages = {190},
	title = {{TOAST: improving reference-free cell composition estimation by cross-cell type differential analysis}},
	url = {https://doi.org/10.1186/s13059-019-1778-0},
	volume = {20},
	year = {2019},
	bdsk-url-1 = {https://doi.org/10.1186/s13059-019-1778-0}}


@article{carter2012,
	abstract = {Tumors vary in their ratio of normal to cancerous cells and in their genomic copy number. Carter et al. describe an analytic method for inferring the purity and ploidy of a tumor sample, enabling longitudinal studies of subclonal mutations and tumor evolution.},
	author = {Carter, Scott L and Cibulskis, Kristian and Helman, Elena and others},
	date = {2012/05/01},
	date-added = {2025-02-04 11:24:28 +0100},
	date-modified = {2025-02-04 11:24:28 +0100},
	doi = {10.1038/nbt.2203},
	id = {Carter2012},
	isbn = {1546-1696},
	journal = {Nature Biotechnology},
	number = {5},
	pages = {413--421},
	title = {{Absolute quantification of somatic DNA alterations in human cancer}},
	url = {https://doi.org/10.1038/nbt.2203},
	volume = {30},
	year = {2012},
	bdsk-url-1 = {https://doi.org/10.1038/nbt.2203}}


@inbook{rumelhart1988,
  author  = {Rumelhart, D E and Hinton, G E and Williams, R J},
  title   = {{Readings in Cognitive Science}},
  chapter = {8: Learning internal representations by error propagation},
  pages   = {399--421},
  year    = {1988},
  publisher = {Elsevier},
}

@article{lecun1989,
    author = {LeCun, Y. and Boser, B. and Denker, J. S. and others},
    title = {{Backpropagation Applied to Handwritten Zip Code Recognition}},
    journal = {Neural Computation},
    volume = {1},
    number = {4},
    pages = {541-551},
    year = {1989},
    month = {12},
    abstract = {The ability of learning networks to generalize can be greatly enhanced by providing constraints from the task domain. This paper demonstrates how such constraints can be integrated into a backpropagation network through the architecture of the network. This approach has been successfully applied to the recognition of handwritten zip code digits provided by the U.S. Postal Service. A single network learns the entire recognition operation, going from the normalized image of the character to the final classification.},
    issn = {0899-7667},
    doi = {10.1162/neco.1989.1.4.541},
    url = {https://doi.org/10.1162/neco.1989.1.4.541},
    eprint = {https://direct.mit.edu/neco/article-pdf/1/4/541/811941/neco.1989.1.4.541.pdf},
}


@article{bourlard1988,
	abstract = {The multilayer perceptron, when working in auto-association mode, is sometimes considered as an interesting candidate to perform data compression or dimensionality reduction of the feature space in information processing applications. The present paper shows that, for auto-association, the nonlinearities of the hidden units are useless and that the optimal parameter values can be derived directly by purely linear techniques relying on singular value decomposition and low rank matrix approximation, similar in spirit to the well-known Karhunen-Lo{\`e}ve transform. This approach appears thus as an efficient alternative to the general error back-propagation algorithm commonly used for training multilayer perceptrons. Moreover, it also gives a clear interpretation of the r{\^o}le of the different parameters.},
	author = {Bourlard, H. and Kamp, Y.},
	date = {1988/09/01},
	date-added = {2025-02-04 11:39:23 +0100},
	date-modified = {2025-02-04 11:39:23 +0100},
	doi = {10.1007/BF00332918},
	id = {Bourlard1988},
	isbn = {1432-0770},
	journal = {Biological Cybernetics},
	number = {4},
	pages = {291--294},
	title = {{Auto-association by multilayer perceptrons and singular value decomposition}},
	url = {https://doi.org/10.1007/BF00332918},
	volume = {59},
	year = {1988},
	bdsk-url-1 = {https://doi.org/10.1007/BF00332918}}


Systematic reviews to support evidence-based medicine, 2nd edition (2nd ed.). CRC Press. https://doi.org/10.1201/b13411

@book{khan2011,
	author = {Khan, K. and Kunz, R. and Kleijnen, J. and others},
	doi = {10.1201/b13411},
	publisher = {CRC Press},
	title = {{Systematic reviews to support evidence-based medicine, 2nd edition}},
	url = {https://doi.org/10.1201/b13411},
	year = {2011},
    address = {London}
}

@article{moher2010,
title = {{Preferred reporting items for systematic reviews and meta-analyses: The PRISMA statement}},
journal = {International Journal of Surgery},
volume = {8},
number = {5},
pages = {336-341},
year = {2010},
issn = {1743-9191},
doi = {https://doi.org/10.1016/j.ijsu.2010.02.007},
url = {https://www.sciencedirect.com/science/article/pii/S1743919110000403},
author = {David Moher and Alessandro Liberati and Jennifer Tetzlaff and others},
keywords = {Reporting guidelines, Systematic reviews, Meta-analyses, Evaluating health interventions, Improving quality}
}

@article{page2021a,
	author = {Page, Matthew J. and McKenzie, Joanne E. and Bossuyt, Patrick M. and others},
	date = {2021/03/29},
	date-added = {2025-02-04 11:44:02 +0100},
	date-modified = {2025-02-04 11:44:02 +0100},
	doi = {10.1186/s13643-021-01626-4},
	id = {Page2021},
	isbn = {2046-4053},
	journal = {Systematic Reviews},
	number = {1},
	pages = {89},
	title = {{The PRISMA 2020 statement: an updated guideline for reporting systematic reviews}},
	url = {https://doi.org/10.1186/s13643-021-01626-4},
	volume = {10},
	year = {2021},
	bdsk-url-1 = {https://doi.org/10.1186/s13643-021-01626-4}}

@article{page2021b,
title = {{Updating guidance for reporting systematic reviews: development of the PRISMA 2020 statement}},
journal = {Journal of Clinical Epidemiology},
volume = {134},
pages = {103-112},
year = {2021},
issn = {0895-4356},
doi = {https://doi.org/10.1016/j.jclinepi.2021.02.003},
url = {https://www.sciencedirect.com/science/article/pii/S0895435621000408},
author = {Matthew J Page and Joanne E McKenzie and Patrick M Bossuyt and others},
keywords = {Systematic reviews, Meta-analysis, Reporting guidelines, Transparency, Reproducibility},
abstract = {Objectives
To describe the processes used to update the PRISMA 2009 statement for reporting systematic reviews, present results of a survey conducted to inform the update, summarize decisions made at the PRISMA update meeting, and describe and justify changes made to the guideline.
Methods
We reviewed 60 documents with reporting guidance for systematic reviews to generate suggested modifications to the PRISMA 2009 statement. We invited 220 systematic review methodologists and journal editors to complete a survey about the suggested modifications. The results of these projects were discussed at a 21-member in-person meeting. Following the meeting, we drafted the PRISMA 2020 statement and refined it based on feedback from co-authors and a convenience sample of 15 systematic reviewers.
Results
The review of 60 documents revealed that all topics addressed by the PRISMA 2009 statement could be modified. Of the 110 survey respondents, more than 66% recommended keeping six of the original checklist items as they were and modifying 15 of them using wording suggested by us. Attendees at the in-person meeting supported the revised wording for several items but suggested rewording for most to enhance clarity, and further refinements were made over six drafts of the guideline.
Conclusions
The PRISMA 2020 statement consists of updated reporting guidance for systematic reviews. We hope that providing this detailed description of the development process will enhance the acceptance and uptake of the guideline and assist those developing and updating future reporting guidelines.}
}

@ARTICLE{torroja2019,
AUTHOR={Torroja, Carlos  and S\'anchez-Cabo, F\'atima },
TITLE={{Digitaldlsorter: Deep-Learning on scRNA-Seq to Deconvolute Gene Expression Data}},
JOURNAL={Frontiers in Genetics},
VOLUME={10},
YEAR={2019},
URL={https://www.frontiersin.org/journals/genetics/articles/10.3389/fgene.2019.00978},
DOI={10.3389/fgene.2019.00978},
ISSN={1664-8021},
ABSTRACT={<p>The development of single cell transcriptome sequencing has allowed researchers the possibility to dig inside the role of the individual cell types in a plethora of disease scenarios. It also expands to the whole transcriptome what before was only possible for a few tenths of antibodies in cell population analysis. More importantly, it allows resolving the permanent question of whether the changes observed in a particular bulk experiment are a consequence of changes in cell type proportions or an aberrant behavior of a particular cell type. However, single cell experiments are still complex to perform and expensive to sequence making bulk RNA-Seq experiments yet more common. scRNA-Seq data is proving highly relevant information for the characterization of the immune cell repertoire in different diseases ranging from cancer to atherosclerosis. In particular, as scRNA-Seq becomes more widely used, new types of immune cell populations emerge and their role in the genesis and evolution of the disease opens new avenues for personalized immune therapies. Immunotherapy have already proven successful in a variety of tumors such as breast, colon and melanoma and its value in other types of disease is being currently explored. From a statistical perspective, single-cell data are particularly interesting due to its high dimensionality, overcoming the limitations of the “skinny matrix” that traditional bulk RNA-Seq experiments yield. With the technological advances that enable sequencing hundreds of thousands of cells, scRNA-Seq data have become especially suitable for the application of Machine Learning algorithms such as Deep Learning (DL). We present here a DL based method to enumerate and quantify the immune infiltration in colorectal and breast cancer bulk RNA-Seq samples starting from scRNA-Seq. Our method makes use of a Deep Neural Network (DNN) model that allows quantification not only of lymphocytes as a general population but also of specific CD8+, CD4Tmem, CD4Th and CD4Tregs subpopulations, as well as B-cells and Stromal content. Moreover, the signatures are built from scRNA-Seq data from the tumor, preserving the specific characteristics of the tumor microenvironment as opposite to other approaches in which cells were isolated from blood. Our method was applied to synthetic bulk RNA-Seq and to samples from the TCGA project yielding very accurate results in terms of quantification and survival prediction.</p>}}


@article{
menden2020,
author = {Kevin Menden and Mohamed Marouf and Sergio Oller and others},
title = {{Deep learning–based cell composition analysis from tissue expression profiles}},
journal = {Science Advances},
volume = {6},
number = {30},
pages = {eaba2619},
year = {2020},
doi = {10.1126/sciadv.aba2619},
URL = {https://www.science.org/doi/abs/10.1126/sciadv.aba2619},
eprint = {https://www.science.org/doi/pdf/10.1126/sciadv.aba2619},
abstract = {Scaden enables robust cell type deconvolution of complex samples across data types, using a deep learning–based model. We present Scaden, a deep neural network for cell deconvolution that uses gene expression information to infer the cellular composition of tissues. Scaden is trained on single-cell RNA sequencing (RNA-seq) data to engineer discriminative features that confer robustness to bias and noise, making complex data preprocessing and feature selection unnecessary. We demonstrate that Scaden outperforms existing deconvolution algorithms in both precision and robustness. A single trained network reliably deconvolves bulk RNA-seq and microarray, human and mouse tissue expression data and leverages the combined information of multiple datasets. Because of this stability and flexibility, we surmise that deep learning will become an algorithmic mainstay for cell deconvolution of various data types. Scaden’s software package and web application are easy to use on new as well as diverse existing expression datasets available in public resources, deepening the molecular and cellular understanding of developmental and disease processes.}}


@Article{liu2022,
AUTHOR = {Liu, Zhendong and Lv, Xinrong and Chen, Xi and others},
TITLE = {{Predicting Algorithm of Tissue Cell Ratio Based on Deep Learning Using Single-Cell RNA Sequencing}},
JOURNAL = {Applied Sciences},
VOLUME = {12},
YEAR = {2022},
NUMBER = {12},
ARTICLE-NUMBER = {5790},
URL = {https://www.mdpi.com/2076-3417/12/12/5790},
ISSN = {2076-3417},
ABSTRACT = {Background: Understanding the proportion of cell types in heterogeneous tissue samples is important in bioinformatics. It is a challenge to infer the proportion of tissues using bulk RNA sequencing data in bioinformatics because most traditional algorithms for predicting tissue cell ratios heavily rely on standardized specific cell-type gene expression profiles, and do not consider tissue heterogeneity. The prediction accuracy of algorithms is limited, and robustness is lacking. This means that new approaches are needed urgently. Methods: In this study, we introduced an algorithm that automatically predicts tissue cell ratios named Autoptcr. The algorithm uses the data simulated by single-cell RNA sequencing (ScRNA-Seq) for model training, using convolutional neural networks (CNNs) to extract intrinsic relationships between genes and predict the cell proportions of tissues. Results: We trained the algorithm using simulated bulk samples and made predictions using real bulk PBMC data. Comparing Autoptcr with existing advanced algorithms, the Pearson correlation coefficient between the actual value of Autoptcr and the predicted value was the highest, reaching 0.903. Tested on a bulk sample, the correlation coefficient of Lin was 41% higher than that of CSx. The algorithm can infer tissue cell proportions directly from tissue gene expression data. Conclusions: The Autoptcr algorithm uses simulated ScRNA-Seq data for training to solve the problem of specific cell-type gene expression profiles. It also has high prediction accuracy and strong noise resistance for the tissue cell ratio. This work is expected to provide new research ideas for the prediction of tissue cell proportions.},
DOI = {10.3390/app12125790}
}

@article{lin2022,
title = {{DAISM-DNNXMBD: Highly accurate cell type proportion estimation with in silico data augmentation and deep neural networks}},
journal = {Patterns},
volume = {3},
number = {3},
pages = {100440},
year = {2022},
issn = {2666-3899},
doi = {https://doi.org/10.1016/j.patter.2022.100440},
url = {https://www.sciencedirect.com/science/article/pii/S2666389922000137},
author = {Yating Lin and Haojun Li and Xu Xiao and others},
keywords = {cell type proportion estimation, deconvolution, data augmentation, data simulation, deep learning},
abstract = {Summary
Understanding the immune cell abundance of cancer and other disease-related tissues has an important role in guiding disease treatments. Computational cell type proportion estimation methods have been previously developed to derive such information from bulk RNA sequencing data. Unfortunately, our results show that the performance of these methods can be seriously plagued by the mismatch between training data and real-world data. To tackle this issue, we propose the DAISM-DNNXMBD (XMBD: Xiamen Big Data, a biomedical open software initiative in the National Institute for Data Science in Health and Medicine, Xiamen University, China.) (denoted as DAISM-DNN) pipeline that trains a deep neural network (DNN) with dataset-specific training data populated from a certain amount of calibrated samples using DAISM, a novel data augmentation method with an in silico mixing strategy. The evaluation results demonstrate that the DAISM-DNN pipeline outperforms other existing methods consistently and substantially for all the cell types under evaluation in real-world datasets.}
}


@article{chen2022,
	abstract = {Single-cell RNA-sequencing has become a powerful tool to study biologically significant characteristics at explicitly high resolution. However, its application on emerging data is currently limited by its intrinsic techniques. Here, we introduce Tissue-AdaPtive autoEncoder (TAPE), a deep learning method connecting bulk RNA-seq and single-cell RNA-seq to achieve precise deconvolution in a short time. By constructing an interpretable decoder and training under a unique scheme, TAPE can predict cell-type fractions and cell-type-specific gene expression tissue-adaptively. Compared with popular methods on several datasets, TAPE has a better overall performance and comparable accuracy at cell type level. Additionally, it is more robust among different cell types, faster, and sensitive to provide biologically meaningful predictions. Moreover, through the analysis of clinical data, TAPE shows its ability to predict cell-type-specific gene expression profiles with biological significance. We believe that TAPE will enable and accelerate the precise analysis of high-throughput clinical data in a wide range.},
	author = {Chen, Yanshuo and Wang, Yixuan and Chen, Yuelong and others},
	date = {2022/11/08},
	date-added = {2025-02-04 11:49:54 +0100},
	date-modified = {2025-02-04 11:49:54 +0100},
	doi = {10.1038/s41467-022-34550-9},
	id = {Chen2022},
	isbn = {2041-1723},
	journal = {Nature Communications},
	number = {1},
	pages = {6735},
	title = {{Deep autoencoder for interpretable tissue-adaptive deconvolution and cell-type-specific gene analysis}},
	url = {https://doi.org/10.1038/s41467-022-34550-9},
	volume = {13},
	year = {2022},
	bdsk-url-1 = {https://doi.org/10.1038/s41467-022-34550-9}}


@article{chiu2023,
	abstract = {Cell composition deconvolution (CCD) is a type of bioinformatic task to estimate the cell fractions from bulk gene expression profiles, such as RNA-seq. Many CCD models were developed to perform linear regression analysis using reference gene expression signatures of distinct cell types. Reference gene expression signatures could be generated from cell-specific gene expression profiles, such as scRNA-seq. However, the batch effects and dropout events frequently observed across scRNA-seq datasets have limited the performances of CCD methods.},
	author = {Chiu, Yen-Jung and Ni, Chung-En and Huang, Yen-Hua},
	date = {2023/10/31},
	date-added = {2025-02-04 11:50:41 +0100},
	date-modified = {2025-02-04 11:50:41 +0100},
	doi = {10.1186/s12920-023-01674-w},
	id = {Chiu2023},
	isbn = {1755-8794},
	journal = {BMC Medical Genomics},
	number = {2},
	pages = {272},
	title = {{HArmonized single-cell RNA-seq Cell type Assisted Deconvolution (HASCAD)}},
	url = {https://doi.org/10.1186/s12920-023-01674-w},
	volume = {16},
	year = {2023},
	bdsk-url-1 = {https://doi.org/10.1186/s12920-023-01674-w}}


@article{charytonowicz2023,
	abstract = {We introduce UniCell: Deconvolve Base (UCDBase), a pre-trained, interpretable, deep learning model to deconvolve cell type fractions and predict cell identity across Spatial, bulk-RNA-Seq, and scRNA-Seq datasets without contextualized reference data. UCD is trained on 10 million pseudo-mixtures from a fully-integrated scRNA-Seq training database comprising over 28 million annotated single cells spanning 840 unique cell types from 898 studies. We show that our UCDBase and transfer-learning models achieve comparable or superior performance on in-silico mixture deconvolution to existing, reference-based, state-of-the-art methods. Feature attribute analysis uncovers gene signatures associated with cell-type specific inflammatory-fibrotic responses in ischemic kidney injury, discerns cancer subtypes, and accurately deconvolves tumor microenvironments. UCD identifies pathologic changes in cell fractions among bulk-RNA-Seq data for several disease states. Applied to lung cancer scRNA-Seq data, UCD annotates and distinguishes normal from cancerous cells. Overall, UCD enhances transcriptomic data analysis, aiding in assessment of cellular and spatial context.},
	author = {Charytonowicz, Daniel and Brody, Rachel and Sebra, Robert},
	date = {2023/03/11},
	date-added = {2025-02-04 11:51:27 +0100},
	date-modified = {2025-02-04 11:51:27 +0100},
	doi = {10.1038/s41467-023-36961-8},
	id = {Charytonowicz2023},
	isbn = {2041-1723},
	journal = {Nature Communications},
	number = {1},
	pages = {1350},
	title = {{Interpretable and context-free deconvolution of multi-scale whole transcriptomic data with UniCell deconvolve}},
	url = {https://doi.org/10.1038/s41467-023-36961-8},
	volume = {14},
	year = {2023},
	bdsk-url-1 = {https://doi.org/10.1038/s41467-023-36961-8}}


@article{yan2023,
	abstract = {Prior research has shown that the deconvolution of cell-free RNA can uncover the tissue origin. The conventional deconvolution approaches rely on constructing a reference tissue-specific gene panel, which cannot capture the inherent variation present in actual data. To address this, we have developed a novel method that utilizes a neural network framework to leverage the entire training dataset. Our approach involved training a model that incorporated 15 distinct tissue types. Through one semi-independent and two complete independent validations, including deconvolution using a semi in silico dataset, deconvolution with a custom normal tissue mixture RNA-seq data, and deconvolution of longitudinal circulating tumor cell RNA-seq (ctcRNA) data from a cancer patient with metastatic tumors, we demonstrate the efficacy and advantages of the deep-learning approach which were exerted by effectively capturing the inherent variability present in the dataset, thus leading to enhanced accuracy. Sensitivity analyses reveal that neural network models are less susceptible to the presence of missing data, making them more suitable for real-world applications. Moreover, by leveraging the concept of organotropism, we applied our approach to trace the migration of circulating tumor cell-derived RNA (ctcRNA) in a cancer patient with metastatic tumors, thereby highlighting the potential clinical significance of early detection of cancer metastasis.},
	author = {Yan, Fengyao and Jiang, Limin and Ye, Fei and others},
	date = {2023/11/04},
	date-added = {2025-02-04 11:53:40 +0100},
	date-modified = {2025-02-04 11:53:40 +0100},
	doi = {10.1186/s12967-023-04663-w},
	id = {Yan2023},
	isbn = {1479-5876},
	journal = {Journal of Translational Medicine},
	number = {1},
	pages = {783},
	title = {{Deep neural network based tissue deconvolution of circulating tumor cell RNA}},
	url = {https://doi.org/10.1186/s12967-023-04663-w},
	volume = {21},
	year = {2023},
	bdsk-url-1 = {https://doi.org/10.1186/s12967-023-04663-w}}


@article{jin2024,
	abstract = {The composition of cell-type is a key indicator of health. Advancements in bulk gene expression data curation, single cell RNA-sequencing technologies, and computational deconvolution approaches offer a new perspective to learn about the composition of different cell types in a quick and affordable way. In this study, we developed a quantile regression and deep learning-based method called Neural Network Immune Contexture Estimator (NNICE) to estimate the cell type abundance and its uncertainty by automatically deconvolving bulk RNA-seq data. The proposed NNICE model was able to successfully recover ground-truth cell type fraction values given unseen bulk mixture gene expression profiles from the same dataset it was trained on. Compared with baseline methods, NNICE achieved better performance on deconvolve both pseudo-bulk gene expressions (Pearson correlation R = 0.9) and real bulk gene expression data (Pearson correlation R = 0.9) across all cell types. In conclusion, NNICE combines statistic inference with deep learning to provide accurate and interpretable cell type deconvolution from bulk gene expression.},
	author = {Jin, Yong Won and Hu, Pingzhao and Liu, Qian},
	date = {2024/06/18},
	date-added = {2025-02-04 11:54:18 +0100},
	date-modified = {2025-02-04 11:54:18 +0100},
	doi = {10.1038/s41598-024-65053-w},
	id = {Jin2024},
	isbn = {2045-2322},
	journal = {Scientific Reports},
	number = {1},
	pages = {14040},
	title = {{NNICE: a deep quantile neural network algorithm for expression deconvolution}},
	url = {https://doi.org/10.1038/s41598-024-65053-w},
	volume = {14},
	year = {2024},
	bdsk-url-1 = {https://doi.org/10.1038/s41598-024-65053-w}}

@article{huang2024,
title = {{DeepDecon accurately estimates cancer cell fractions in bulk RNA-seq data}},
journal = {Patterns},
volume = {5},
number = {5},
pages = {100969},
year = {2024},
issn = {2666-3899},
doi = {https://doi.org/10.1016/j.patter.2024.100969},
url = {https://www.sciencedirect.com/science/article/pii/S2666389924000771},
author = {Jiawei Huang and Yuxuan Du and Andres Stucky and others},
keywords = {deconvolution, scRNA-seq, deep neural network, cancer tissue},
abstract = {Summary
Understanding the cellular composition of a disease-related tissue is important in disease diagnosis, prognosis, and downstream treatment. Recent advances in single-cell RNA-sequencing (scRNA-seq) technique have allowed the measurement of gene expression profiles for individual cells. However, scRNA-seq is still too expensive to be used for large-scale population studies, and bulk RNA-seq is still widely used in such situations. An essential challenge is to deconvolve cellular composition for bulk RNA-seq data based on scRNA-seq data. Here, we present DeepDecon, a deep neural network model that leverages single-cell gene expression information to accurately predict the fraction of cancer cells in bulk tissues. It provides a refining strategy in which the cancer cell fraction is iteratively estimated by a set of trained models. When applied to simulated and real cancer data, DeepDecon exhibits superior performance compared to existing decomposition methods in terms of accuracy.}
}


@article{khatri2024,
	abstract = {Cell deconvolution is the estimation of cell type fractions and cell type-specific gene expression from mixed data. An unmet challenge in cell deconvolution is the scarcity of realistic training data and the domain shift often observed in synthetic training data. Here, we show that two novel deep neural networks with simultaneous consistency regularization of the target and training domains significantly improve deconvolution performance. Our algorithm, DISSECT, outperforms competing algorithms in cell fraction and gene expression estimation by up to 14 percentage points. DISSECT can be easily adapted to other biomedical data types, as exemplified by our proteomic deconvolution experiments.},
	author = {Khatri, Robin and Machart, Pierre and Bonn, Stefan},
	date = {2024/04/30},
	date-added = {2025-02-04 12:06:09 +0100},
	date-modified = {2025-02-04 12:06:09 +0100},
	doi = {10.1186/s13059-024-03251-5},
	id = {Khatri2024},
	isbn = {1474-760X},
	journal = {Genome Biology},
	number = {1},
	pages = {112},
	title = {{DISSECT: deep semi-supervised consistency regularization for accurate cell type fraction and gene expression estimation}},
	url = {https://doi.org/10.1186/s13059-024-03251-5},
	volume = {25},
	year = {2024},
	bdsk-url-1 = {https://doi.org/10.1186/s13059-024-03251-5}}

@article{english2019,
author = {Andrew English and Nigel Irwin},
title ={{Nonclassical Islet Peptides: Pancreatic and Extrapancreatic Actions}},
journal = {Clinical Medicine Insights: Endocrinology and Diabetes},
volume = {12},
number = {},
pages = {1179551419888871},
year = {2019},
doi = {10.1177/1179551419888871},
note ={PMID: 32425629},
URL = {https://doi.org/10.1177/1179551419888871},
eprint = {https://doi.org/10.1177/1179551419888871}, 
abstract = { The pancreas has physiologically important endocrine and exocrine functions; secreting enzymes into the small intestine to aid digestion and releasing multiple peptide hormones via the islets of Langerhans to regulate glucose metabolism, respectively. Insulin and glucagon, in combination with ghrelin, pancreatic polypeptide and somatostatin, are the main classical islet peptides critical for the maintenance of blood glucose. However, pancreatic islets also synthesis numerous ‘nonclassical’ peptides that have recently been demonstrated to exert fundamental effects on overall islet function and metabolism. As such, insights into the physiological relevance of these nonclassical peptides have shown impact on glucose metabolism, insulin action, cell survival, weight loss, and energy expenditure. This review will focus on the role of individual nonclassical islet peptides to stimulate pancreatic islet secretions as well as regulate metabolism. In addition, the more recognised actions of these peptides on satiety and energy regulation will also be considered. Furthermore, recent advances in the field of peptide therapeutics and obesity-diabetes have focused on the benefits of simultaneously targeting several hormone receptor signalling cascades. The potential for nonclassical islet hormones within such combinational approaches will also be discussed. }
}


@article{newman2019,
	abstract = {Single-cell RNA-sequencing has emerged as a powerful technique for characterizing cellular heterogeneity, but it is currently impractical on large sample cohorts and cannot be applied to fixed specimens collected as part of routine clinical care. We previously developed an approach for digital cytometry, called CIBERSORT, that enables estimation of cell type abundances from bulk tissue transcriptomes. We now introduce CIBERSORTx, a machine learning method that extends this framework to infer cell-type-specific gene expression profiles without physical cell isolation. By minimizing platform-specific variation, CIBERSORTx also allows the use of single-cell RNA-sequencing data for large-scale tissue dissection. We evaluated the utility of CIBERSORTx in multiple tumor types, including melanoma, where single-cell reference profiles were used to dissect bulk clinical specimens, revealing cell-type-specific phenotypic states linked to distinct driver mutations and response to immune checkpoint blockade. We anticipate that digital cytometry will augment single-cell profiling efforts, enabling cost-effective, high-throughput tissue characterization without the need for antibodies, disaggregation or viable cells.},
	author = {Newman, Aaron M. and Steen, Chlo{\'e} B. and Liu, Chih Long and others},
	date = {2019/07/01},
	date-added = {2025-02-04 12:10:02 +0100},
	date-modified = {2025-02-04 12:10:02 +0100},
	doi = {10.1038/s41587-019-0114-2},
	id = {Newman2019},
	isbn = {1546-1696},
	journal = {Nature Biotechnology},
	number = {7},
	pages = {773--782},
	title = {{Determining cell type abundance and expression from bulk tissues with digital cytometry}},
	url = {https://doi.org/10.1038/s41587-019-0114-2},
	volume = {37},
	year = {2019},
	bdsk-url-1 = {https://doi.org/10.1038/s41587-019-0114-2}}


@article{finotello2019,
	abstract = {It was highlighted that the original article {$[$}1{$]$} contained a typesetting mistake in the name of Noel Filipe da Cunha Carvalho de Miranda. This was incorrectly captured as Noel Filipe da Cunha Carvahlo de Miranda. It was also highlighted that in Fig. 3C the left panels Y-axis were cropped and in Fig. 5C, CD8 bar was cropped. This Correction article shows the correct Figs. 3 and 5. The original article has been updated.},
	author = {Finotello, Francesca and Mayer, Clemens and Plattner, Christina and others},
	date = {2019/07/29},
	date-added = {2025-02-04 12:10:43 +0100},
	date-modified = {2025-02-04 12:10:43 +0100},
	doi = {10.1186/s13073-019-0655-5},
	id = {Finotello2019},
	isbn = {1756-994X},
	journal = {Genome Medicine},
	number = {1},
	pages = {50},
	title = {{Correction to: Molecular and pharmacological modulators of the tumor immune contexture revealed by deconvolution of RNA-seq data}},
	url = {https://doi.org/10.1186/s13073-019-0655-5},
	volume = {11},
	year = {2019},
	bdsk-url-1 = {https://doi.org/10.1186/s13073-019-0655-5}}


@article{becht2016,
	abstract = {We introduce the Microenvironment Cell Populations-counter (MCP-counter) method, which allows the robust quantification of the absolute abundance of eight immune and two stromal cell populations in heterogeneous tissues from transcriptomic data. We present in vitro mRNA mixture and ex vivo immunohistochemical data that quantitatively support the validity of our method's estimates. Additionally, we demonstrate that MCP-counter overcomes several limitations or weaknesses of previously proposed computational approaches. MCP-counter is applied to draw a global picture of immune infiltrates across human healthy tissues and non-hematopoietic human tumors and recapitulates microenvironment-based patient stratifications associated with overall survival in lung adenocarcinoma and colorectal and breast cancer.},
	author = {Becht, Etienne and Giraldo, Nicolas A. and Lacroix, Laetitia and others},
	date = {2016/10/20},
	date-added = {2025-02-04 12:11:29 +0100},
	date-modified = {2025-02-04 12:11:29 +0100},
	doi = {10.1186/s13059-016-1070-5},
	id = {Becht2016},
	isbn = {1474-760X},
	journal = {Genome Biology},
	number = {1},
	pages = {218},
	title = {{Estimating the population abundance of tissue-infiltrating immune and stromal cell populations using gene expression}},
	url = {https://doi.org/10.1186/s13059-016-1070-5},
	volume = {17},
	year = {2016},
	bdsk-url-1 = {https://doi.org/10.1186/s13059-016-1070-5}}


@article{frishberg2019,
	abstract = {Single-cell RNA sequencing (scRNA-seq) is a rich resource of cellular heterogeneity, opening new avenues in the study of complex tissues. We introduce Cell Population Mapping (CPM), a deconvolution algorithm in which reference scRNA-seq profiles are leveraged to infer the composition of cell types and states from bulk transcriptome data (`scBio'CRAN R-package). Analysis of individual variations in lungs of influenza-virus-infected mice reveals that the relationship between cell abundance and clinical symptoms is a cell-state-specific property that varies gradually along the continuum of cell-activation states. The gradual change is confirmed in subsequent experiments and is further explained by a mathematical model in which clinical outcomes relate to cell-state dynamics along the activation process. Our results demonstrate the power of CPM in reconstructing the continuous spectrum of cell states within heterogeneous tissues.},
	author = {Frishberg, Amit and Peshes-Yaloz, Naama and Cohn, Ofir and others},
	date = {2019/04/01},
	date-added = {2025-02-04 12:12:03 +0100},
	date-modified = {2025-02-04 12:12:03 +0100},
	doi = {10.1038/s41592-019-0355-5},
	id = {Frishberg2019},
	isbn = {1548-7105},
	journal = {Nature Methods},
	number = {4},
	pages = {327--332},
	title = {{Cell composition analysis of bulk genomics using single-cell data}},
	url = {https://doi.org/10.1038/s41592-019-0355-5},
	volume = {16},
	year = {2019},
	bdsk-url-1 = {https://doi.org/10.1038/s41592-019-0355-5}}

@article{li2017,
    author = {Li, Taiwen and Fan, Jingyu and Wang, Binbin and others},
    title = {{TIMER: A Web Server for Comprehensive Analysis of Tumor-Infiltrating Immune Cells}},
    journal = {Cancer Research},
    volume = {77},
    number = {21},
    pages = {e108-e110},
    year = {2017},
    month = {10},
    abstract = {Recent clinical successes of cancer immunotherapy necessitate the investigation of the interaction between malignant cells and the host immune system. However, elucidation of complex tumor–immune interactions presents major computational and experimental challenges. Here, we present Tumor Immune Estimation Resource (TIMER; cistrome.shinyapps.io/timer) to comprehensively investigate molecular characterization of tumor–immune interactions. Levels of six tumor-infiltrating immune subsets are precalculated for 10,897 tumors from 32 cancer types. TIMER provides 6 major analytic modules that allow users to interactively explore the associations between immune infiltrates and a wide spectrum of factors, including gene expression, clinical outcomes, somatic mutations, and somatic copy number alterations. TIMER provides a user-friendly web interface for dynamic analysis and visualization of these associations, which will be of broad utilities to cancer researchers. Cancer Res; 77(21); e108–10. ©2017 AACR.},
    issn = {0008-5472},
    doi = {10.1158/0008-5472.CAN-17-0307},
    url = {https://doi.org/10.1158/0008-5472.CAN-17-0307},
    eprint = {https://aacrjournals.org/cancerres/article-pdf/77/21/e108/2934090/e108.pdf},
}

@article{monaco2019,
title = {{RNA-Seq Signatures Normalized by mRNA Abundance Allow Absolute Deconvolution of Human Immune Cell Types}},
journal = {Cell Reports},
volume = {26},
number = {6},
pages = {1627-1640.e7},
year = {2019},
issn = {2211-1247},
doi = {https://doi.org/10.1016/j.celrep.2019.01.041},
url = {https://www.sciencedirect.com/science/article/pii/S2211124719300592},
author = {Gianni Monaco and Bernett Lee and Weili Xu and others},
keywords = {immune system, flow cytometry, transcriptome, RNA-seq, gene modules, housekeeping, mRNA composition, mRNA abundance, mRNA heterogeneity, deconvolution},
abstract = {Summary
The molecular characterization of immune subsets is important for designing effective strategies to understand and treat diseases. We characterized 29 immune cell types within the peripheral blood mononuclear cell (PBMC) fraction of healthy donors using RNA-seq (RNA sequencing) and flow cytometry. Our dataset was used, first, to identify sets of genes that are specific, are co-expressed, and have housekeeping roles across the 29 cell types. Then, we examined differences in mRNA heterogeneity and mRNA abundance revealing cell type specificity. Last, we performed absolute deconvolution on a suitable set of immune cell types using transcriptomics signatures normalized by mRNA abundance. Absolute deconvolution is ready to use for PBMC transcriptomic data using our Shiny app (https://github.com/giannimonaco/ABIS). We benchmarked different deconvolution and normalization methods and validated the resources in independent cohorts. Our work has research, clinical, and diagnostic value by making it possible to effectively associate observations in bulk transcriptomics data to specific immune subsets.}
}

@article{erdmann-pham2021,
author = {Erdmann-Pham, Dan D. and Fischer, Jonathan and Hong, Justin and others}, 
title = {{Likelihood-based deconvolution of bulk gene expression data using single-cell references}},
volume = {31}, 
number = {10}, 
pages = {1794-1806}, 
year = {2021}, 
doi = {10.1101/gr.272344.120}, 
abstract ={Direct comparison of bulk gene expression profiles is complicated by distinct cell type mixtures in each sample that obscure whether observed differences are actually caused by changes in the expression levels themselves or are simply a result of differing cell type compositions. Single-cell technology has made it possible to measure gene expression in individual cells, achieving higher resolution at the expense of increased noise. If carefully incorporated, such single-cell data can be used to deconvolve bulk samples to yield accurate estimates of the true cell type proportions, thus enabling one to disentangle the effects of differential expression and cell type mixtures. Here, we propose a generative model and a likelihood-based inference method that uses asymptotic statistical theory and a novel optimization procedure to perform deconvolution of bulk RNA-seq data to produce accurate cell type proportion estimates. We show the effectiveness of our method, called RNA-Sieve, across a diverse array of scenarios involving real data and discuss extensions made uniquely possible by our probabilistic framework, including a demonstration of well-calibrated confidence intervals.}, 
URL = {http://genome.cshlp.org/content/31/10/1794.abstract}, 
eprint = {http://genome.cshlp.org/content/31/10/1794.full.pdf+html}, 
journal = {Genome Research} 
}


@article{tsoucas2019,
	abstract = {The rapid development of single-cell transcriptomic technologies has helped uncover the cellular heterogeneity within cell populations. However, bulk RNA-seq continues to be the main workhorse for quantifying gene expression levels due to technical simplicity and low cost. To most effectively extract information from bulk data given the new knowledge gained from single-cell methods, we have developed a novel algorithm to estimate the cell-type composition of bulk data from a single-cell RNA-seq-derived cell-type signature. Comparison with existing methods using various real RNA-seq data sets indicates that our new approach is more accurate and comprehensive than previous methods, especially for the estimation of rare cell types. More importantly, our method can detect cell-type composition changes in response to external perturbations, thereby providing a valuable, cost-effective method for dissecting the cell-type-specific effects of drug treatments or condition changes. As such, our method is applicable to a wide range of biological and clinical investigations.},
	author = {Tsoucas, Daphne and Dong, Rui and Chen, Haide and others},
	date = {2019/07/05},
	date-added = {2025-02-04 12:16:50 +0100},
	date-modified = {2025-02-04 12:16:50 +0100},
	doi = {10.1038/s41467-019-10802-z},
	id = {Tsoucas2019},
	isbn = {2041-1723},
	journal = {Nature Communications},
	number = {1},
	pages = {2975},
	title = {{Accurate estimation of cell-type composition from gene expression data}},
	url = {https://doi.org/10.1038/s41467-019-10802-z},
	volume = {10},
	year = {2019},
	bdsk-url-1 = {https://doi.org/10.1038/s41467-019-10802-z}}


@article{jew2020,
	abstract = {We present Bisque, a tool for estimating cell type proportions in bulk expression. Bisque implements a regression-based approach that utilizes single-cell RNA-seq (scRNA-seq) or single-nucleus RNA-seq (snRNA-seq) data to generate a reference expression profile and learn gene-specific bulk expression transformations to robustly decompose RNA-seq data. These transformations significantly improve decomposition performance compared to existing methods when there is significant technical variation in the generation of the reference profile and observed bulk expression. Importantly, compared to existing methods, our approach is extremely efficient, making it suitable for the analysis of large genomic datasets that are becoming ubiquitous. When applied to subcutaneous adipose and dorsolateral prefrontal cortex expression datasets with both bulk RNA-seq and snRNA-seq data, Bisque replicates previously reported associations between cell type proportions and measured phenotypes across abundant and rare cell types. We further propose an additional mode of operation that merely requires a set of known marker genes.},
	author = {Jew, Brandon and Alvarez, Marcus and Rahmani, Elior and others},
	date = {2020/04/24},
	date-added = {2025-02-04 12:17:29 +0100},
	date-modified = {2025-02-04 12:17:29 +0100},
	doi = {10.1038/s41467-020-15816-6},
	id = {Jew2020},
	isbn = {2041-1723},
	journal = {Nature Communications},
	number = {1},
	pages = {1971},
	title = {{Accurate estimation of cell composition in bulk expression through robust integration of single-cell information}},
	url = {https://doi.org/10.1038/s41467-020-15816-6},
	volume = {11},
	year = {2020},
	bdsk-url-1 = {https://doi.org/10.1038/s41467-020-15816-6}}

@misc{xie2024,
      title={{Statistical Inference for Cell Type Deconvolution}}, 
      author={Dongyue Xie and Jingshu Wang},
      year={2024},
      eprint={2202.06420},
      archivePrefix={arXiv},
      primaryClass={stat.ME},
      url={https://arxiv.org/abs/2202.06420}, 
}

@article{dong2020,
    author = {Dong, Meichen and Thennavan, Aatish and Urrutia, Eugene and others},
    title = {{SCDC: bulk gene expression deconvolution by multiple single-cell RNA sequencing references}},
    journal = {Briefings in Bioinformatics},
    volume = {22},
    number = {1},
    pages = {416-427},
    year = {2020},
    month = {01},
    abstract = {Recent advances in single-cell RNA sequencing (scRNA-seq) enable characterization of transcriptomic profiles with single-cell resolution and circumvent averaging artifacts associated with traditional bulk RNA sequencing (RNA-seq) data. Here, we propose SCDC, a deconvolution method for bulk RNA-seq that leverages cell-type specific gene expression profiles from multiple scRNA-seq reference datasets. SCDC adopts an ENSEMBLE method to integrate deconvolution results from different scRNA-seq datasets that are produced in different laboratories and at different times, implicitly addressing the problem of batch-effect confounding. SCDC is benchmarked against existing methods using both in silico generated pseudo-bulk samples and experimentally mixed cell lines, whose known cell-type compositions serve as ground truths. We show that SCDC outperforms existing methods with improved accuracy of cell-type decomposition under both settings. To illustrate how the ENSEMBLE framework performs in complex tissues under different scenarios, we further apply our method to a human pancreatic islet dataset and a mouse mammary gland dataset. SCDC returns results that are more consistent with experimental designs and that reproduce more significant associations between cell-type proportions and measured phenotypes.},
    issn = {1477-4054},
    doi = {10.1093/bib/bbz166},
    url = {https://doi.org/10.1093/bib/bbz166},
    eprint = {https://academic.oup.com/bib/article-pdf/22/1/416/35934954/bbz166.pdf},
}


@article{chu2022,
	abstract = {Inferring single-cell compositions and their contributions to global gene expression changes from bulk RNA sequencing (RNA-seq) datasets is a major challenge in oncology. Here we develop Bayesian cell proportion reconstruction inferred using statistical marginalization (BayesPrism), a Bayesian method to predict cellular composition and gene expression in individual cell types from bulk RNA-seq, using patient-derived, scRNA-seq as prior information. We conduct integrative analyses in primary glioblastoma, head and neck squamous cell carcinoma and skin cutaneous melanoma to correlate cell type composition with clinical outcomes across tumor types, and explore spatial heterogeneity in malignant and nonmalignant cell states. We refine current cancer subtypes using gene expression annotation after exclusion of confounding nonmalignant cells. Finally, we identify genes whose expression in malignant cells correlates with macrophage infiltration, T cells, fibroblasts and endothelial cells across multiple tumor types. Our work introduces a new lens to accurately infer cellular composition and expression in large cohorts of bulk RNA-seq data.},
	author = {Chu, Tinyi and Wang, Zhong and Pe'er, Dana and others},
	date = {2022/04/01},
	date-added = {2025-02-04 12:22:29 +0100},
	date-modified = {2025-02-04 12:22:29 +0100},
	doi = {10.1038/s43018-022-00356-3},
	id = {Chu2022},
	isbn = {2662-1347},
	journal = {Nature Cancer},
	number = {4},
	pages = {505--517},
	title = {{Cell type and gene expression deconvolution with BayesPrism enables Bayesian integrative analysis across bulk and single-cell RNA sequencing in oncology}},
	url = {https://doi.org/10.1038/s43018-022-00356-3},
	volume = {3},
	year = {2022},
	bdsk-url-1 = {https://doi.org/10.1038/s43018-022-00356-3}}

@article{wang2021,
author = {Wang, Jiebiao and Roeder, Kathryn and Devlin, Bernie}, 
title = {{Bayesian estimation of cell type–specific gene expression with prior derived from single-cell data}},
volume = {31}, 
number = {10}, 
pages = {1807-1818}, 
year = {2021}, 
doi = {10.1101/gr.268722.120}, 
abstract ={When assessed over a large number of samples, bulk RNA sequencing provides reliable data for gene expression at the tissue level. Single-cell RNA sequencing (scRNA-seq) deepens those analyses by evaluating gene expression at the cellular level. Both data types lend insights into disease etiology. With current technologies, scRNA-seq data are known to be noisy. Constrained by costs, scRNA-seq data are typically generated from a relatively small number of subjects, which limits their utility for some analyses, such as identification of gene expression quantitative trait loci (eQTLs). To address these issues while maintaining the unique advantages of each data type, we develop a Bayesian method (bMIND) to integrate bulk and scRNA-seq data. With a prior derived from scRNA-seq data, we propose to estimate sample-level cell type–specific (CTS) expression from bulk expression data. The CTS expression enables large-scale sample-level downstream analyses, such as detection of CTS differentially expressed genes (DEGs) and eQTLs. Through simulations, we show that bMIND improves the accuracy of sample-level CTS expression estimates and increases the power to discover CTS DEGs when compared to existing methods. To further our understanding of two complex phenotypes, autism spectrum disorder and Alzheimer's disease, we apply bMIND to gene expression data of relevant brain tissue to identify CTS DEGs. Our results complement findings for CTS DEGs obtained from snRNA-seq studies, replicating certain DEGs in specific cell types while nominating other novel genes for those cell types. Finally, we calculate CTS eQTLs for 11 brain regions by analyzing Genotype-Tissue Expression Project data, creating a new resource for biological insights.}, 
URL = {http://genome.cshlp.org/content/31/10/1807.abstract}, 
eprint = {http://genome.cshlp.org/content/31/10/1807.full.pdf+html}, 
journal = {Genome Research} 
}

@inproceedings{goodfellow2014,
 author = {Goodfellow, Ian and Pouget-Abadie, Jean and Mirza, Mehdi and others},
 booktitle = {Advances in Neural Information Processing Systems},
 pages = {2672 - 2680},
 publisher = {Curran Associates, Inc.},
 title = {{Generative Adversarial Nets}},
 url = {https://proceedings.neurips.cc/paper_files/paper/2014/file/5ca3e9b122f61f8f06494c97b1afccf3-Paper.pdf},
 volume = {2},
 year = {2014}
}

@article{thomas2025,
	title = {{GBMPurity}: {A} {Machine} {Learning} {Tool} for {Estimating} {Glioblastoma} {Tumour} {Purity} from {Bulk} {RNA}-seq {Data}},
	issn = {1522-8517},
	url = {https://doi.org/10.1093/neuonc/noaf026},
	doi = {10.1093/neuonc/noaf026},
	abstract = {Glioblastoma (GBM) presents a significant clinical challenge due to its aggressive nature and extensive heterogeneity. Tumour purity, the proportion of malignant cells within a tumour, is an important covariate for understanding the disease, having direct clinical relevance or obscuring signal of the malignant portion in molecular analyses of bulk samples. However, current methods for estimating tumour purity are non-specific and technically demanding. Therefore, we aimed to build a reliable and accessible purity estimator for GBM.We developed GBMPurity, a deep-learning model specifically designed to estimate the purity of IDH-wildtype primary GBM from bulk RNA-seq data. The model was trained using simulated pseudobulk tumours of known purity from labelled single-cell data acquired from the GBmap resource. The performance of GBMPurity was evaluated and compared to several existing tools using independent datasets.GBMPurity outperformed existing tools, achieving a mean absolute error of 0.15 and a concordance correlation coefficient of 0.88 on validation datasets. We demonstrate the utility of GBMPurity through inference on bulk RNA-seq samples and observe reduced purity of the Proneural molecular subtype relative to the Classical, attributed to the increased presence of healthy brain cells.GBMPurity provides a reliable and accessible tool for estimating tumour purity from bulk RNA-seq data, enhancing the interpretation of bulk RNA-seq data and offering valuable insights into GBM biology. To facilitate the use of this model by the wider research community, GBMPurity is available as a web-based tool at: https://gbmdeconvoluter.leeds.ac.uk/.},
	journal = {Neuro-Oncology},
	author = {Thomas, Morgan P H and Ajaib, Shoaib and Tanner, Georgette and others},
	month = feb,
	year = {2025},
	note = {\_eprint: https://academic.oup.com/neuro-oncology/advance-article-pdf/doi/10.1093/neuonc/noaf026/61724213/noaf026.pdf},
	pages = {noaf026},
}

@article{sun2024,
title = {{Deciphering the abundance of immune cells in glomerular endothelium of Alport syndrome kidneys using the deconvolution algorithm CONVdeconv}},
journal = {Future Generation Computer Systems},
volume = {161},
pages = {496-501},
year = {2024},
issn = {0167-739X},
doi = {https://doi.org/10.1016/j.future.2024.07.013},
url = {https://www.sciencedirect.com/science/article/pii/S0167739X24003765},
author = {Yizhou Sun and Jiao Liu and Fang Fang and others},
keywords = {Neural network, Alport syndrome, Pseudo bulk, Single-cell RNA},
abstract = {Due to the high cost of single-cell sequencing technology, understanding cell heterogeneity within tissues is crucial for elucidating the biological characteristics of complex tissues. Therefore, this study proposes a method for generating pseudo-bulk data based on single-cell sequencing data and designs a convolutional neural network model, CONVdeconv, based on attention mechanisms, for deconvoluting real bulk RNA data. Experimental validation on mouse kidney data demonstrates that the CONVdeconv model performs exceptionally well in deconvolution tasks, effectively restoring cell type proportions. Furthermore, deconvolution analysis of mouse data with Alport syndrome revealed significant differences in macrophages and T lymphocytes between disease and normal states, providing new clues and directions for a deeper understanding of the disease’s progression.}
}

@Article{D’Agostino2022,
author={D'Agostino, Nunzio
and Li, Wenli
and Wang, Dapeng},
title={{High-throughput transcriptomics}},
journal={Scientific Reports},
year={2022},
month={Nov},
day={29},
volume={12},
number={1},
pages={20313},
abstract={High-throughput transcriptomics has revolutionised the field of transcriptome research by offering a cost-effective and powerful screening tool. Standard bulk RNA sequencing (RNA-Seq) enables characterisation of the average expression profiles for individual samples and facilitates identification of the molecular functions associated with genes differentially expressed across conditions. RNA-Seq can also be applied to disentangle splicing variants and discover novel transcripts, thus contributing to a comprehensive understanding of the transcriptome landscape. A closely related technique, single-cell RNA-Seq, has enabled the study of cell-type-specific gene expressions in hundreds to thousands of cells, aiding the exploration of cell heterogeneity. Nowadays, bulk RNA-Seq and single-cell RNA-Seq serve as complementary tools to advance and accelerate the development of transcriptome-based resources. This Collection illustrates how the current global research community makes use of these techniques to address a broad range of questions in life sciences. It demonstrates the usefulness and popularity of high-throughput transcriptomics and presents the best practices and potential issues for the benefit of future end-users.},
issn={2045-2322},
doi={10.1038/s41598-022-23985-1},
url={https://doi.org/10.1038/s41598-022-23985-1}
}

@book{Rumelhart1986,
author = {Rumelhart, David E. and Mcclelland, James L.},
publisher= {MIT Press},
year = {1986},
month = {01},
address = {Cambridge, Massachusetts},
title = {{Parallel distributed processing: explorations in the microstructure of cognition. Volume 1. Foundations}}
}

@article{heydari2022,
    author = {Heydari, A. Ali and Davalos, Oscar A. and Zhao, Lihong and others},
    title = {{ACTIVA: realistic single-cell RNA-seq generation with automatic cell-type identification using introspective variational autoencoders}},
    journal = {Bioinformatics},
    volume = {38},
    number = {8},
    pages = {2194-2201},
    year = {2022},
    month = {02},
    abstract = {Single-cell RNA sequencing (scRNAseq) technologies allow for measurements of gene expression at a single-cell resolution. This provides researchers with a tremendous advantage for detecting heterogeneity, delineating cellular maps or identifying rare subpopulations. However, a critical complication remains: the low number of single-cell observations due to limitations by rarity of subpopulation, tissue degradation or cost. This absence of sufficient data may cause inaccuracy or irreproducibility of downstream analysis. In this work, we present Automated Cell-Type-informed Introspective Variational Autoencoder (ACTIVA): a novel framework for generating realistic synthetic data using a single-stream adversarial variational autoencoder conditioned with cell-type information. Within a single framework, ACTIVA can enlarge existing datasets and generate specific subpopulations on demand, as opposed to two separate models [such as single-cell GAN (scGAN) and conditional scGAN (cscGAN)]. Data generation and augmentation with ACTIVA can enhance scRNAseq pipelines and analysis, such as benchmarking new algorithms, studying the accuracy of classifiers and detecting marker genes. ACTIVA will facilitate analysis of smaller datasets, potentially reducing the number of patients and animals necessary in initial studies.We train and evaluate models on multiple public scRNAseq datasets. In comparison to GAN-based models (scGAN and cscGAN), we demonstrate that ACTIVA generates cells that are more realistic and harder for classifiers to identify as synthetic which also have better pair-wise correlation between genes. Data augmentation with ACTIVA significantly improves classification of rare subtypes (more than 45\% improvement compared with not augmenting and 4\% better than cscGAN) all while reducing run-time by an order of magnitude in comparison to both models.The codes and datasets are hosted on Zenodo (https://doi.org/10.5281/zenodo.5879639). Tutorials are available at https://github.com/SindiLab/ACTIVA.Supplementary data are available at Bioinformatics online.},
    issn = {1367-4803},
    doi = {10.1093/bioinformatics/btac095},
    url = {https://doi.org/10.1093/bioinformatics/btac095},
    eprint = {https://academic.oup.com/bioinformatics/article-pdf/38/8/2194/49010211/btac095.pdf},
}

@Article{Marouf2020,
author={Marouf, Mohamed
and Machart, Pierre
and Bansal, Vikas
and others},
title={{Realistic in silico generation and augmentation of single-cell RNA-seq data using generative adversarial networks}},
journal={Nature Communications},
year={2020},
month={Jan},
day={09},
volume={11},
number={1},
pages={166},
abstract={A fundamental problem in biomedical research is the low number of observations available, mostly due to a lack of available biosamples, prohibitive costs, or ethical reasons. Augmenting few real observations with generated in silico samples could lead to more robust analysis results and a higher reproducibility rate. Here, we propose the use of conditional single-cell generative adversarial neural networks (cscGAN) for the realistic generation of single-cell RNA-seq data. cscGAN learns non-linear gene--gene dependencies from complex, multiple cell type samples and uses this information to generate realistic cells of defined types. Augmenting sparse cell populations with cscGAN generated cells improves downstream analyses such as the detection of marker genes, the robustness and reliability of classifiers, the assessment of novel analysis algorithms, and might reduce the number of animal experiments and costs in consequence. cscGAN outperforms existing methods for single-cell RNA-seq data generation in quality and hold great promise for the realistic generation and augmentation of other biomedical data types.},
issn={2041-1723},
doi={10.1038/s41467-019-14018-z},
url={https://doi.org/10.1038/s41467-019-14018-z}
}

@Article{Treppner2021,
author={Treppner, Martin
and Salas-Bastos, Adri{\'a}n
and Hess, Moritz
and others},
title={{Synthetic single cell RNA sequencing data from small pilot studies using deep generative models}},
journal={Scientific Reports},
year={2021},
month={Apr},
day={30},
volume={11},
number={1},
pages={9403},
abstract={Deep generative models, such as variational autoencoders (VAEs) or deep Boltzmann machines (DBMs), can generate an arbitrary number of synthetic observations after being trained on an initial set of samples. This has mainly been investigated for imaging data but could also be useful for single-cell transcriptomics (scRNA-seq). A small pilot study could be used for planning a full-scale experiment by investigating planned analysis strategies on synthetic data with different sample sizes. It is unclear whether synthetic observations generated based on a small scRNA-seq dataset reflect the properties relevant for subsequent data analysis steps. We specifically investigated two deep generative modeling approaches, VAEs and DBMs. First, we considered single-cell variational inference (scVI) in two variants, generating samples from the posterior distribution, the standard approach, or the prior distribution. Second, we propose single-cell deep Boltzmann machines (scDBMs). When considering the similarity of clustering results on synthetic data to ground-truth clustering, we find that the {\$}{\$}scVI{\_}{\{}posterior{\}}{\$}{\$}variant resulted in high variability, most likely due to amplifying artifacts of small datasets. All approaches showed mixed results for cell types with different abundance by overrepresenting highly abundant cell types and missing less abundant cell types. With increasing pilot dataset sizes, the proportions of the cells in each cluster became more similar to that of ground-truth data. We also showed that all approaches learn the univariate distribution of most genes, but problems occurred with bimodality. Across all analyses, in comparing 10{\$}{\$}{\backslash}times{\$}{\$}Genomics and Smart-seq2 technologies, we could show that for 10{\$}{\$}{\backslash}times{\$}{\$}datasets, which have higher sparsity, it is more challenging to make inference from small to larger datasets. Overall, the results show that generative deep learning approaches might be valuable for supporting the design of scRNA-seq experiments.},
issn={2045-2322},
doi={10.1038/s41598-021-88875-4},
url={https://doi.org/10.1038/s41598-021-88875-4}
}

﻿@Article{Chung2017,
author={Chung, Woosung
and Eum, Hye Hyeon
and Lee, Hae-Ock
and others},
title={{Single-cell RNA-seq enables comprehensive tumour and immune cell profiling in primary breast cancer}},
journal={Nature Communications},
year={2017},
month={May},
day={05},
volume={8},
number={1},
pages={15081},
abstract={Single-cell transcriptome profiling of tumour tissue isolates allows the characterization of heterogeneous tumour cells along with neighbouring stromal and immune cells. Here we adopt this powerful approach to breast cancer and analyse 515 cells from 11 patients. Inferred copy number variations from the single-cell RNA-seq data separate carcinoma cells from non-cancer cells. At a single-cell resolution, carcinoma cells display common signatures within the tumour as well as intratumoral heterogeneity regarding breast cancer subtype and crucial cancer-related pathways. Most of the non-cancer cells are immune cells, with three distinct clusters of T lymphocytes, B lymphocytes and macrophages. T lymphocytes and macrophages both display immunosuppressive characteristics: T cells with a regulatory or an exhausted phenotype and macrophages with an M2 phenotype. These results illustrate that the breast cancer transcriptome has a wide range of intratumoral heterogeneity, which is shaped by the tumour cells and immune cells in the surrounding microenvironment.},
issn={2041-1723},
doi={10.1038/ncomms15081},
url={https://doi.org/10.1038/ncomms15081}
}

@article{Nogales2021,
title = {{A survey of deep learning models in medical therapeutic areas}},
journal = {Artificial Intelligence in Medicine},
volume = {112},
pages = {102020},
year = {2021},
issn = {0933-3657},
doi = {https://doi.org/10.1016/j.artmed.2021.102020},
url = {https://www.sciencedirect.com/science/article/pii/S0933365721000130},
author = {Alberto Nogales and Álvaro J. García-Tejedor and Diana Monge and others},
keywords = {Survey, Artificial intelligence, Deep learning, Medicine},
abstract = {Artificial intelligence is a broad field that comprises a wide range of techniques, where deep learning is presently the one with the most impact. Moreover, the medical field is an area where data both complex and massive and the importance of the decisions made by doctors make it one of the fields in which deep learning techniques can have the greatest impact. A systematic review following the Cochrane recommendations with a multidisciplinary team comprised of physicians, research methodologists and computer scientists has been conducted. This survey aims to identify the main therapeutic areas and the deep learning models used for diagnosis and treatment tasks. The most relevant databases included were MedLine, Embase, Cochrane Central, Astrophysics Data System, Europe PubMed Central, Web of Science and Science Direct. An inclusion and exclusion criteria were defined and applied in the first and second peer review screening. A set of quality criteria was developed to select the papers obtained after the second screening. Finally, 126 studies from the initial 3493 papers were selected and 64 were described. Results show that the number of publications on deep learning in medicine is increasing every year. Also, convolutional neural networks are the most widely used models and the most developed area is oncology where they are used mainly for image analysis.}
}

@Article{Salman2024,
author={Salman, Diaa
and Direkoglu, Cem
and Kusaf, Mehmet
and others},
title={{Hybrid deep learning models for time series forecasting of solar power}},
journal={Neural Computing and Applications},
year={2024},
month={Jun},
day={01},
volume={36},
number={16},
pages={9095-9112},
abstract={Forecasting solar power production accurately is critical for effectively planning and managing renewable energy systems. This paper introduces and investigates novel hybrid deep learning models for solar power forecasting using time series data. The research analyzes the efficacy of various models for capturing the complex patterns present in solar power data. In this study, all of the possible combinations of convolutional neural network (CNN), long short-term memory (LSTM), and transformer (TF) models are experimented. These hybrid models also compared with the single CNN, LSTM and TF models with respect to different kinds of optimizers. Three different evaluation metrics are also employed for performance analysis. Results show that the CNN--LSTM--TF hybrid model outperforms the other models, with a mean absolute error (MAE) of 0.551{\%} when using the Nadam optimizer. However, the TF--LSTM model has relatively low performance, with an MAE of 16.17{\%}, highlighting the difficulties in making reliable predictions of solar power. This result provides valuable insights for optimizing and planning renewable energy systems, highlighting the significance of selecting appropriate models and optimizers for accurate solar power forecasting. This is the first time such a comprehensive work presented that also involves transformer networks in hybrid models for solar power forecasting.},
issn={1433-3058},
doi={10.1007/s00521-024-09558-5},
url={https://doi.org/10.1007/s00521-024-09558-5}
}

@article{Bayoudh2024,
title = {{A survey of multimodal hybrid deep learning for computer vision: Architectures, applications, trends, and challenges}},
journal = {Information Fusion},
volume = {105},
pages = {102217},
year = {2024},
issn = {1566-2535},
doi = {https://doi.org/10.1016/j.inffus.2023.102217},
url = {https://www.sciencedirect.com/science/article/pii/S156625352300533X},
author = {Khaled Bayoudh},
keywords = {Applications, Computer vision, Multimodal hybrid deep learning, Sensory modalities},
abstract = {In recent years, deep learning algorithms have rapidly revolutionized artificial intelligence, particularly machine learning, enabling researchers and practitioners to extend previously hand-crafted feature extraction procedures. In particular, deep learning uses adaptive learning processes to learn more complex and informative patterns from datasets of varying sizes. With the increasing availability of multimodal data streams and recent advances in deep learning algorithms, multimodal deep learning is on the rise. This requires the development of complex models that can process and analyze multimodal information in a consistent manner. However, unstructured data can come in many different forms (also known as modalities). Extracting relevant features from this data remains an ambitious goal for deep learning researchers. According to the literature, most deep learning systems consist of a single architecture (i.e., standalone deep learning). When two or more deep learning architectures are combined over multiple sensory modalities, the result is called a multimodal hybrid deep learning model. Since this research direction has received much attention in the field of deep learning, the purpose of this survey is to provide a broader overview of the topic. In this paper, we provide a comprehensive review of recent advances in multimodal hybrid deep learning, including a thorough analysis of the most commonly developed hybrid architectures. In particular, one of the main challenges in multimodal hybrid analysis is the ability of these architectures to systematically integrate cross-modal features in hybrid designs. Therefore, we propose a generic framework for multimodal hybrid learning that focuses mainly on fusion methods. We also identify trends and challenges in multimodal hybrid learning and provide insights and directions for future research. Our findings show that multimodal hybrid learning can perform well in a variety of challenging computer vision applications and tasks.}
}

@Article{Mavaie2023,
author={Mavaie, Pegah
and Holder, Lawrence
and Skinner, Michael K.},
title={{Hybrid deep learning approach to improve classification of low-volume high-dimensional data}},
journal={BMC Bioinformatics},
year={2023},
month={Nov},
day={07},
volume={24},
number={1},
pages={419},
abstract={The performance of machine learning classification methods relies heavily on the choice of features. In many domains, feature generation can be labor-intensive and require domain knowledge, and feature selection methods do not scale well in high-dimensional datasets. Deep learning has shown success in feature generation but requires large datasets to achieve high classification accuracy. Biology domains typically exhibit these challenges with numerous handcrafted features (high-dimensional) and small amounts of training data (low volume).},
issn={1471-2105},
doi={10.1186/s12859-023-05557-w},
url={https://doi.org/10.1186/s12859-023-05557-w}
}

@article{Pasquini2021,
title = {{Automated methods for cell type annotation on scRNA-seq data}},
journal = {Computational and Structural Biotechnology Journal},
volume = {19},
pages = {961-969},
year = {2021},
issn = {2001-0370},
doi = {https://doi.org/10.1016/j.csbj.2021.01.015},
url = {https://www.sciencedirect.com/science/article/pii/S2001037021000192},
author = {Giovanni Pasquini and Jesus Eduardo {Rojo Arias} and Patrick Schäfer and others},
keywords = {scRNA-seq, Cell type, Cell state, Automatic annotation},
abstract = {The advent of single-cell sequencing started a new era of transcriptomic and genomic research, advancing our knowledge of the cellular heterogeneity and dynamics. Cell type annotation is a crucial step in analyzing single-cell RNA sequencing data, yet manual annotation is time-consuming and partially subjective. As an alternative, tools have been developed for automatic cell type identification. Different strategies have emerged to ultimately associate gene expression profiles of single cells with a cell type either by using curated marker gene databases, correlating reference expression data, or transferring labels by supervised classification. In this review, we present an overview of the available tools and the underlying approaches to perform automated cell type annotations on scRNA-seq data.}
}

@ARTICLE{Sen2018,

AUTHOR={Sen, Partho  and Kemppainen, Esko  and Orešič, Matej},
TITLE={{Perspectives on Systems Modeling of Human Peripheral Blood Mononuclear Cells}},
JOURNAL={Frontiers in Molecular Biosciences},
VOLUME={4},
YEAR={2018},
URL={https://www.frontiersin.org/journals/molecular-biosciences/articles/10.3389/fmolb.2017.00096},
DOI={10.3389/fmolb.2017.00096},
ISSN={2296-889X},
ABSTRACT={<p>Human peripheral blood mononuclear cells (PBMCs) are the key drivers of the immune responses. These cells undergo activation, proliferation and differentiation into various subsets. During these processes they initiate metabolic reprogramming, which is coordinated by specific gene and protein activities. PBMCs as a model system have been widely used to study metabolic and autoimmune diseases. Herein we review various omics and systems-based approaches such as transcriptomics, epigenomics, proteomics, and metabolomics as applied to PBMCs, particularly T helper subsets, that unveiled disease markers and the underlying mechanisms. We also discuss and emphasize several aspects of T cell metabolic modeling in healthy and disease states using genome-scale metabolic models.</p>}}

@Article{Lopez2022,
author={Lopez, Romain
and Li, Baoguo
and Keren-Shaul, Hadas
and others},
title={{DestVI identifies continuums of cell types in spatial transcriptomics data}},
journal={Nature Biotechnology},
year={2022},
month={Sep},
day={01},
volume={40},
number={9},
pages={1360-1369},
abstract={Most spatial transcriptomics technologies are limited by their resolution, with spot sizes larger than that of a single cell. Although joint analysis with single-cell RNA sequencing can alleviate this problem, current methods are limited to assessing discrete cell types, revealing the proportion of cell types inside each spot. To identify continuous variation of the transcriptome within cells of the same type, we developed Deconvolution of Spatial Transcriptomics profiles using Variational Inference (DestVI). Using simulations, we demonstrate that DestVI outperforms existing methods for estimating gene expression for every cell type inside every spot. Applied to a study of infected lymph nodes and of a mouse tumor model, DestVI provides high-resolution, accurate spatial characterization of the cellular organization of these tissues and identifies cell-type-specific changes in gene expression between different tissue regions or between conditions. DestVI is available as part of the open-source software package scvi-tools (https://scvi-tools.org).},
issn={1546-1696},
doi={10.1038/s41587-022-01272-8},
url={https://doi.org/10.1038/s41587-022-01272-8}
}

@Article{Revkov2023,
author={Revkov, Egor
and Kulshrestha, Tanmay
and Sung, Ken Wing-Kin
and others},
title={{PUREE: accurate pan-cancer tumor purity estimation from gene expression data}},
journal={Communications Biology},
year={2023},
month={Apr},
day={11},
volume={6},
number={1},
pages={394},
abstract={Tumors are complex masses composed of malignant and non-malignant cells. Variation in tumor purity (proportion of cancer cells in a sample) can both confound integrative analysis and enable studies of tumor heterogeneity. Here we developed PUREE, which uses a weakly supervised learning approach to infer tumor purity from a tumor gene expression profile. PUREE was trained on gene expression data and genomic consensus purity estimates from 7864 solid tumor samples. PUREE predicted purity with high accuracy across distinct solid tumor types and generalized to tumor samples from unseen tumor types and cohorts. Gene features of PUREE were further validated using single-cell RNA-seq data from distinct tumor types. In a comprehensive benchmark, PUREE outperformed existing transcriptome-based purity estimation approaches. Overall, PUREE is a highly accurate and versatile method for estimating tumor purity and interrogating tumor heterogeneity from bulk tumor gene expression data, which can complement genomics-based approaches or be used in settings where genomic data is unavailable.},
issn={2399-3642},
doi={10.1038/s42003-023-04764-8},
url={https://doi.org/10.1038/s42003-023-04764-8}
}

@article{zhou2025,
title = {{AGR, LMR and SIRI are the optimal combinations for risk stratification in advanced patients with non-small cell lung cancer following immune checkpoint blockers}},
journal = {International Immunopharmacology},
volume = {149},
pages = {114215},
year = {2025},
issn = {1567-5769},
doi = {https://doi.org/10.1016/j.intimp.2025.114215},
url = {https://www.sciencedirect.com/science/article/pii/S156757692500205X},
author = {Yun Zhou and Xia Liu and Biwen Wu and others},
keywords = {Non-small cell lung cancer, Immune checkpoint blocker, Risk stratification, Biomarker, Prognosis},
abstract = {Due to drug resistance, a majority of patients with non-small cell lung cancer (NSCLC) experience disease progression following immunotherapy. Therefore, there is an urgent need to develop novel biomarkers to predict the prognosis of NSCLC patients. Clinical data from 544 patients with advanced NSCLC who underwent immune checkpoint blockers (ICBs) at our clinical center were collected in this study. The results indicated that low Albumin-Globulin Ratio (AGR) and Lymphocyte-Monocyte Ratio (LMR) and high Systemic Immune-Inflammation Index (SIRI) were significantly correlated with both poor overall survival (OS) and progression-free survival (PFS) in NSCLC patients (P < 0.01). These three indicators collectively formed the most effective combined model for predicting the prognosis of NSCLC. Importantly, risk stratification based on AGR, LMR and SIRI was better than that based on the TNM stage, and served as an independent predictor of OS and PFS. Notably, the nomogram model developed by risk stratification, sex, age, smoking history, and pathological type demonstrated a good ability to predict the 1 to 5-year OS rates for NSCLC patients. In summary, AGR, LMR, and SIRI represented the optimal combined models for forecasting the prognosis of patients with advanced NSCLC who underwent ICBs, offering promising potential as biomarkers to direct personalized clinical interventions.}
}
